# Retention of DLK1 in the endoplasmic reticulum identifies roles for EGF domain-specific O-glycans in the secretory pathway

**DOI:** 10.1101/2024.08.31.610613

**Authors:** Yuko Tashima, Yohei Tsukamoto, Natsumi Tsukamoto, Yuji Kondo, Ehsan Uddin, Wakako Furukawa, Shiori Go, Hideyuki Takeuchi, Tetsuya Okajima

## Abstract

In the endoplasmic reticulum (ER), O-glycosylation by O-fucose, O-glucose, and O-GlcNAc occurs in the epidermal growth factor-like (EGF) domains of secreted or transmembrane glycoproteins. Previous studies focusing on Notch receptors have revealed the pivotal role of these O-glycans in the cell surface expression of Notch or secretion of truncated Notch fragments. Although it has been demonstrated that O-fucose, O-glucose, and O-GlcNAc stabilize individual EGF domains, their role in the secretory pathway after the completion of the folding process remains unexplored. In this study, we used delta-like 1 homolog (DLK1) containing six consecutive EGF domains as a model glycoprotein to investigate the role of EGF domain-specific O-glycans in the secretory pathway. Semi-quantitative site-specific glycoproteomics of recombinantly expressed DLK1 revealed multiple O-fucose and O-glucose modifications in addition to an unusual EOGT-dependent O-hexose modification. Consistent with the results of the secretion assay, inactivation of the glycosyltransferases modifying O-fucose and O-glucose, but not the newly identified O-hexose, perturbed the transport of DLK1 from the ER during retention using the selective hooks (RUSH) system. Importantly, the absence of O-fucose did not result in an apparent loss of O-glucose modification within the same EGF domain, and vice versa. Given that protein O-fucosyltransferase 1 and protein O-glucosyltransferase 1 activities depend on the folded state of the EGF domains, O-glycans affected DLK1 transport independently of the folding process required for O-glycosylation in the ER. These findings highlight the distinct roles of O-glycans in facilitating the transport of DLK1 from the ER to the cell surface.

## Introduction

Delta-like 1 homolog (DLK1), also known as Pref-1 (Preadipocyte Factor-1), controls the fate of adipocyte progenitors by inhibiting adipogenesis (1, 2). The biological significance of DLK1 has been analyzed in various contexts including genomic imprinting (3), adipogenesis (4), glucose metabolism (5), and cancer (6). Aberrant DLK1 expression in humans is associated with obesity (7) and cancer progression and metastasis (8, 9). Furthermore, DLK1 loss-of-function mutations have been identified in children with familial central precocious puberty, thus suggesting a role in pubertal timing (10, 11). Although the physiological and clinical roles of DLK1 have been well characterized, the molecular mechanisms underlying its action of DLK1 remain elusive. Various target proteins have been suggested to mediate the function of DLK1 (12) including NOTCH receptors (13) (14) (15) and insulin-like growth factor-1 (16). DLK1 also interacts with prohibition 1 (PHB1) and PHB2 via its cytoplasmic domain to regulate mitochondrial function (17).

DLK1 is structurally related to Notch receptors that potentially interact with DLK1 (18–21). The extracellular domains of DLK1 are largely composed of six tandem repeats of an epidermal growth factor-like (EGF) domain followed by a short region modified by mucin-type O-glycans. The EGF domains are subjected to O-glycosylation, including O-Fuc, O-Glc, and O-GlcNAc that modifies Ser or Thr residues at defined positions within the EGF domains. The first implication of the presence of these O-glycans in DLK1 EGF domains was obtained by Edman degradation sequencing of human DLK1 purified from amniotic fluid, and this failed to sequence amino acid residues subjected to post-translational modifications (22). Subsequent studies combining amino acid sequencing by Edman degradation and MALDI-MS analysis similarly indicated the presence of O-linked monosaccharide modifications in mouse DLK1 EGF domains (23). However, the resolution and accuracy achieved by MALDI-MS in previous studies were insufficient to determine the identity of monosaccharide modifications. In addition to O-glycosylation in the EGF domains, elongated O-GalNAc glycans susceptible to sialidase and O-glycosidase digestion have been detected in the extracellular juxtamembrane regions, and Ser228 and Ser237 were initially assigned as modification sites (22). Later, these modification sites were amended to Thr233 and Thr246 (23). More detailed glycan analysis was performed for the mouse DLK1 homolog, suggesting the modification with NeuNAcα2-3Galβl-3(NeuNAcα2-6)GalNAc O-glycans at Thr235, Thr244, Thr248 sites of mouse DLK1, where the former two sites correspond to human Thr233 and Thr246 (23). However, these O-GalNAc glycans are highly heterogeneous; therefore, no definitive information is available for the O-glycans, including EGF-domain-specific O-glycans on human and mouse DLK1, in previous literature or the current protein database.

Currently, comprehensive analyses of O-glycans on multiple EGF domains have been extensively conducted using Notch receptors whose protein expression and function depend on these O-glycans (24–27). The EGF domain contains six conserved cysteine residues that form three disulfide bonds: the first and third cysteines (C^1^-C^3^), C^2^-C^4^, and C^5^-C^6^ (Figure 1A). Enzymes responsible for O-glycosylation include protein O-fucosyltransferase 1 (POFUT1), protein O-glucosyltransferase 1 (POGLUT1), POGLUT2/3, and EGF domain-specific O-linked N-acetylglucosamine transferase (EOGT), all of which are endoplasmic reticulum (ER) luminal proteins. POFUT1 catalyzes O-fucosylation of Ser or Thr residues in the consensus sequence 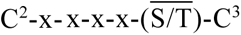(28). POGLUT1 converts O-Glc to a serine in the C^1^-x-S-x-(P/A)-C^2^ (29). EOGT catalyzes the addition of O-GlcNAc (30). O-GlcNAc modifications have been detected in the C^5^-x-x-G-x-(S/T)-G-x-x-C^6^ sequence (31); however, the consensus sequence has not been precisely defined (32, 33). These putative consensus sequences have been used to predict potential glycoproteins modified by O-glycans (31). Importantly, the enzymatic activity of these ER-resident glycosyltransferases requires the folded state of EGF domains, and unfolded peptides are poorly modified by O-glycans *in vitro*.

**Figure 1.**
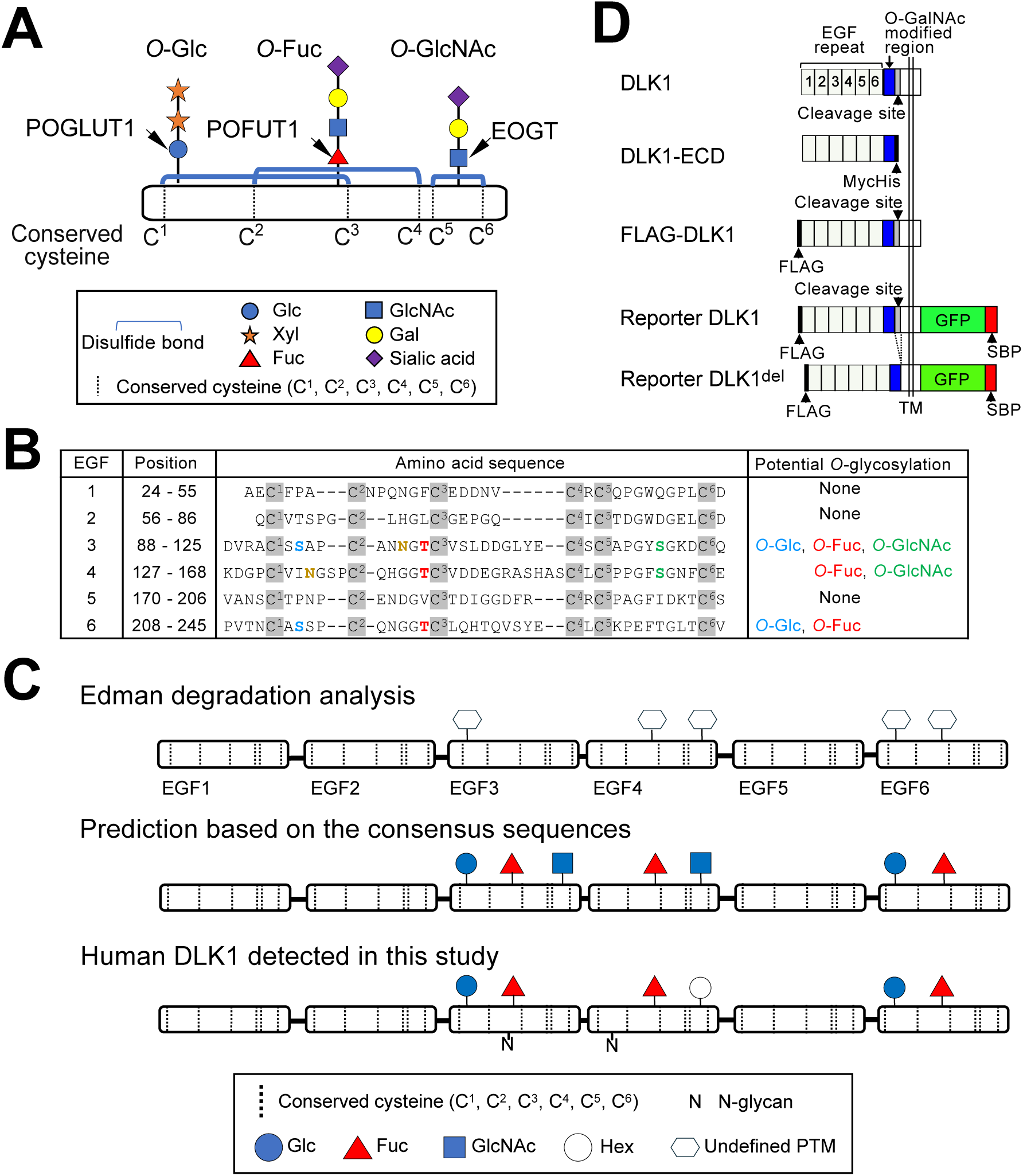
Human DLK1 constructs and O-glycans analyzed in this study. (**A**) EGF domain-specific O-glycans and corresponding O-glycosyltransferases analyzed in this study. The most extended O-glycan structures are presented. Six conserved cysteine residues (C^1^∼C^6^) form three disulfide bonds (*blue*) in a specific pattern. (**B**) The amino acid sequence of six consecutive EGF domains of DLK1. Potential O-glycosylation and N-glycosylation motifs are indicated. The colored letters indicate predicted sites to be modified with the O-Glc (*blue*), O-Fuc (*red*), O-GlcNAc (*green*), and N-glycan (*brown*). The consensus sequences adopted for *in silico* analysis are C^1^-x-S-x-(P/A)-C^2^ for O-glucosylation, C^2^-x-x-x-x-(S/T)-C^3^ for O-fucosylation, C^5^-x-x-G-x-(S/T)-G-x-x-C^6^ for O-GlcNAcylation, and N-X-T for N-glycosylation. (**C**) Previous and current studies on the EGF domain-specific O-glycosylations of human DLK1. There is an apparent discrepancy between the predicted O-glycosylation sites based on the consensus sequence (31) and experimentally identified posttranslational modifications (PTM) of DLK1 isolated from human amniotic fluid (22). The modifications of DLK1 detected in this study are shown at the bottom. (**D**) Full-length and ectodomain (ECD) constructs of human DLK1 used in this study. TM, transmembrane; SBP, streptavidin binding peptide. The DLK1^del^ construct lacks 23 amino acids (Glu279-Glu301) that include the cleavage sites (arrowhead) recognized by TACE/ADAM17.

O-Glycans on EGF domains mediate protein functions on the cell surface, as represented by their critical roles in Notch-ligand interactions. Furthermore, extensive studies examining Notch receptors have revealed the pivotal role of these O-glycans in the cell surface expression of Notch and the secretion of truncated Notch fragments (34–38). The expression of NOTCH1, NOTCH2, and NOTCH3 on the surface of mouse *Pofut1* null embryonic stem cells was similar to that in wild-type cells (39). *Pofut1* deficiency in mouse hematopoietic stem cells caused a slight decrease in the surface expression of NOTCH1 and NOTCH2, but caused no change in NOTCH3 (40). *Pofut1* knockout Chinese hamster ovary cells exhibit slightly decreased NOTCH1 cell surface expression (41). In HEK293T cells, the lack of *POFUT1* or *POGLUT1* causes severe or mild decreases in cell surface NOTCH1 expression, respectively (34–38). The deficient of *EOGT* did not alter the cell surface expression of NOTCH1 (42). An *in vitro* unfolding experiment revealed that O-glycans stabilize folded EGF domains via intramolecular interactions that facilitate folding in the ER (34) (43).

Although O-glycans may simultaneously affect the transport of the folded EGF domain from the ER, their requirement in the secretory pathway has not yet been directly evaluated. However, exogenously expressed NOTCH receptors are prone to accumulation in the ER (44), making them unsuitable for transport analysis from the ER. Instead, we tested whether DLK1 that contains a smaller number of EGF domains, serves as a model system for evaluating the requirement of O-glycans for cell-surface transport from the ER. First, we determined the repertoire of O-glycans displayed on the EGF domains of human DLK1 ectodomains (ECD) expressed in HEK293T cells and the relative abundances of the respective glycoforms. Based on the O-glycan profile, we revealed that the lack of either O-Glc or O-Fuc glycans, due to the inactivation of *POGLUT1* or *POFUT1*, respectively, resulted in impaired secretion of DLK1-ECD. In parallel with this observation, the cell surface transport of DLK1 was markedly perturbed in mutant cells lacking *POGLUT1* or *POFUT1*. The folding process of the EGF domains was sufficient to allow for the complete modification of the remaining O-glycans. Collectively, our data suggest that the absence of O-glycans affects the transport of DLK1 from the ER rather than the folding process required for O-glycosylation in the ER.

## Results

### Analysis of O-glycosylation on DLK1

A previous study based on Edman degradation sequencing of amniotic fluid DLK1 suggested the presence of five post-translational modifications (PTMs) with Ser or Thr residues (22). These included amino acids at the O-Glc modification site of EGF3, O-Fuc, and O-GlcNAc modification sites of EGF4 and the O-Glc and O-Fuc modification sites of EGF6 (Figure 1B). However, the experimentally identified PTMs were inconsistent with those predicted from the consensus sequences of these modifications (Figure 1C) (31). We also reanalyzed the human urine mass spectrometry dataset from the PeptideAtlas database (Experiment ID 8983, PXD016433) (45), and this resulted in the identification of O-Fuc modifications at EGF4, but not at other sites. In this dataset, trypsin was used for peptide preparation, and this was not optimal for generating glycopeptides suitable for O-glycan analysis. In this study, double digestion with trypsin and chymotrypsin was employed for semi-quantitative site-specific O-glycan analysis of DLK1 EGF domains to elucidate the identity and stoichiometry of PTMs.

To isolate and purify the ectodomain (ECD) of DLK1, DLK1-ECD was expressed in HEK293T cells (Figure 1D) and affinity-purified from the culture medium. Protease digests were obtained using trypsin and chymotrypsin and subjected to liquid chromatography-mass spectrometry (LC-MS) using high-energy collision-induced dissociation (HCD) (Table 1 and Figure 2). The extracted ion chromatogram (EIC) of the detectable glycopeptides revealed that DLK1 EGF4 containing the O-Fuc site was modified with deoxy-hexose (dHex), and EGF6 fragments containing both O-Glc and O-Fuc sites were modified with hexose (Hex) and dHex, suggesting the presence of O-glycans, as predicted in previous studies.

**Table 1.** List of proteolytic fragments of DLK1-ECD containing putative glycosylation sites for EGF domain-specific O-glycosylation.

| Position | Peptides sequence | Glycan<br>moieties | Deamidati<br>on<br>(N-glycan) | Proposed<br>O-glycans | z | Theoretical<br>mass (m/z) | Observed<br>mass (m/z)<br>(Exp.1) | Retention<br>time<br>(min)<br>(Exp.1) | Peak<br>height<br>(Exp. 1) | Ratio (%) |  |  |  |
| --- | --- | --- | --- | --- | --- | --- | --- | --- | --- | --- | --- | --- | --- |
|  |  |  |  |  |  |  |  |  |  | Exp. 1 | Exp. 2 | Exp. 3 | Average |
| 91-111 | AC<sup>1</sup>SSAPC<sup>2</sup>ANNGTC<sup>3</sup>VSLDDGLY | - | + | - | 2 | 1116.946 | 1116.946 | 31.36 | 1.3E+06 | 7.6 | 8.4 | 7.2 | 7.7 |
|  |  | Hex [1] | + | O-Glc | 2 | 1197.972 | 1197.973 | 31.11 | 1.6E+07 | 92.4 | 91.6 | 92.8 | 92.3 |
| 112-122 | ECSCAPGYSGK |  | - |  | 2 | 608.242 | 608.242 | 23.64 | 1.5E+05 | 100 | 100 | 100 | 100.0 |
| 128-150 | DGPC<sup>1</sup>VINGSPC<sup>2</sup>QHGGTC<sup>3</sup>VDDEGR | Fuc [1] | + | O-Fuc | 3 | 878.688 | 878.688 | 26.86 | 1.2E+06 | 15.9 | 7.0 | n.d. | 11.5 |
|  |  | Fuc [1] | - | O-Fuc | 3 | 878.360 | 878.360 | 26.77 | 6.2E+06 | 84.1 | 93 | 100 | 92.4 |
| 151-166 | ASHASC<sup>5</sup>LC<sup>6</sup>PPGFSGNF | - | - | - | 2 | 854.864 | 854.864 | 31.18 | 1.1E+06 | 35.1 | 34.3 | 41.0 | 36.8 |
|  |  | Hex [1] | - | O-Hex | 2 | 935.890 | 935.890 | 30.72 | 2.0E+06 | 64.9 | 65.7 | 59.0 | 63.2 |
| 208-231 | PVTNC<sup>1</sup>ASSPC<sup>2</sup>QNGGTC<sup>3</sup>LQHTQVSY | Hex [1] Fuc [1] | - | O-Glc / O-Fuc | 3 | 992.088 | 992.089 | 27.70 | 1.4E+05 | 100 | 100 | 100 | 100.0 |

2 C<sup>1</sup>–C<sup>6</sup>: six conserved cysteine residues. O-glycan structures have been proposed based on experimental data, consensus sequences for O-glycan 3 modification, and the current knowledge of O-glycan structures.

**Figure 2.**
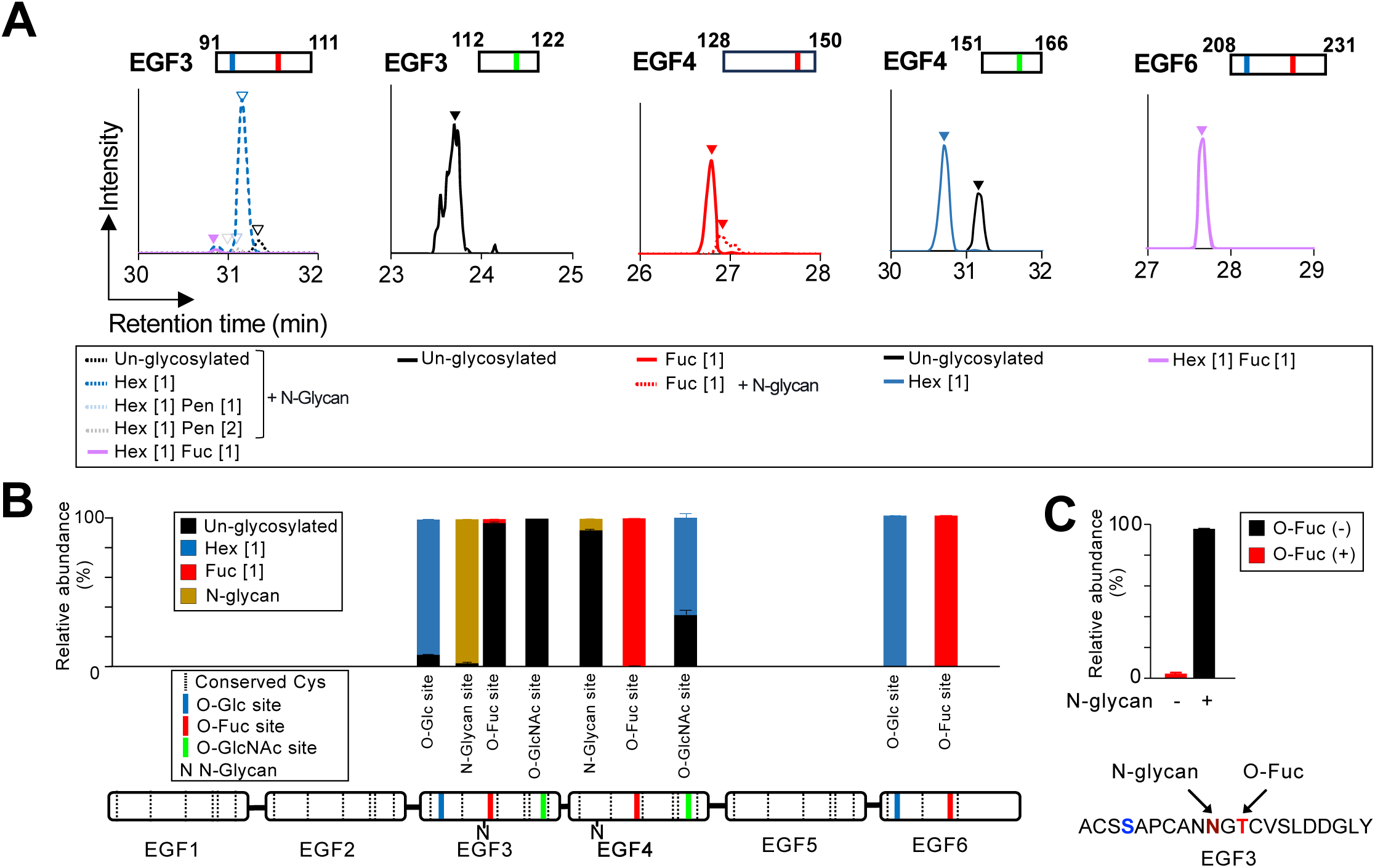
Mass spectrometric analysis of O-glycan modifications on DLK1. DLK1-ECD was purified from cell culture medium of HEK293T cells and digested with trypsin and chymotrypsin. Proteolytic fragments were treated with PNGase F and analyzed using LC-MS/MS. **(A)** Extracted ion chromatograms display unglycosylated and glycosylated peptides that are represented by black and colored lines, respectively. The arrowheads indicate the positions of the precursor ions in the annotated MS/MS spectra (Fig. S2–4). The glycopeptides detected by mass spectrometry are summarized in Table 1. **(B)** Semiquantitative analysis of the glycoforms that modify DLK1 expression. The relative abundance of each glycoform was calculated from the integrated peak height values and presented as the mean ± the range of data (n=3). A schematic of the DLK1 domain structure with predicted glycan modification sites is presented at the bottom. **(C)** Relative abundance of DLK1^[91-111]^ peptides modified with O-fucose and/or N-glycan. Data are presented as the mean ± standard deviation (n=3). The amino acid sequences and modification sites for N-glycans (*brown*) and O-fucose (*red*) are presented.

In contrast, the DLK1 EGF3 fragment containing O-Fuc and O-Glc modification sites predominantly underwent Hex modification, thus indicating modification with O-Glc but not with O-Fuc. Furthermore, the DLK1 EGF3 fragment containing putative O-GlcNAc modification sites was exclusively unglycosylated, although the modification site is conserved in mammals (Figure S1). Notably, HEK293T cells retain endogenous O-GlcNAc transferase activity to generate O-GlcNAc glycans in the NOTCH1 EGF domain (32). These observations are in agreement with data from Edman degradation sequencing (22) but conflict with the predicted O-glycosylation based on consensus sequences.

Two predicted N-glycosylation sites were identified in the human DLK1 cells (Figure 1B). Our analysis detected N-glycosylation predominantly in EGF3 and rarely in EGF4 (Figure 2B). Interestingly, N-glycosylation and O-fucosylation occurred in a mutually exclusive manner in DLK1 EGF3 (Figure 2C). O-fucosylation of EGF3 was detected in only a small population of peptides devoid of N-glycosylation in the N̅GT̅C^3^ sequence. EGF3 with N-glycosylation was barely O-fucosylated, suggesting a potential interplay between N-glycosylation and O-fucosylation in the overlapping consensus sequence.

Two O-GalNAc modifications (Table S1) were detected at Thr256 (Figure S7) and Thr269 (Figure S5 and S8). This is consistent with previously unpublished observations describing these two residues as sites for O-GalNAc modification (23).

Surprisingly, our analysis revealed that the DLK1 EGF4 fragment containing the putative O-GlcNAcylation site was partially modified by Hex (∼60%). This peptide does not include the classical O-Glc modification site or the recently identified site (C^3^-x-N-T-x-G-S-F/Y-x-C^4^) modified with POGLUT2 or POGLUT3 (46, 47). Thus, these data suggest the presence of a novel PTM-modifying EGF domain.

### O-hexose modification of EGF4 depends on EOGT

We further investigated the modification sites and biochemical basis of the newly identified O-Hex modifications. The DLK1 EGF4 sequence 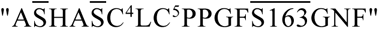 contains three serine residues, including Ser163 at the O-GlcNAcylation site that should be taken into consideration as potential modification sites (Figure 2). Our first attempt at HCD-assisted electron transfer dissociation (EThcD) fragmentation failed to generate fragment ions with glycan moieties, except for mucin-type O-glycans (Figure S5). We investigated if O-Hex modification depends on EOGT activity. To this end, DLK1-ECD was expressed in *EOGT*-deficient HEK293T cells and purified proteins were subjected to LC-MS/MS analysis. EICs corresponding to wild-type and *EOGT*-deficient cells lacked Hex expression in the absence of EOGT (Figure 3). To demonstrate that Ser163 in the EOGT modification site is modified by Hex, we generated a DLK1-ECD^S163A^ mutant. Similar to the absence of *EOGT*, the Hex modification was abolished by the S163A mutation (Figure 3). These results indicate that O-Hex modification occurred at the O-GlcNAc modification site in an *EOGT*-dependent manner.

**Figure 3.**
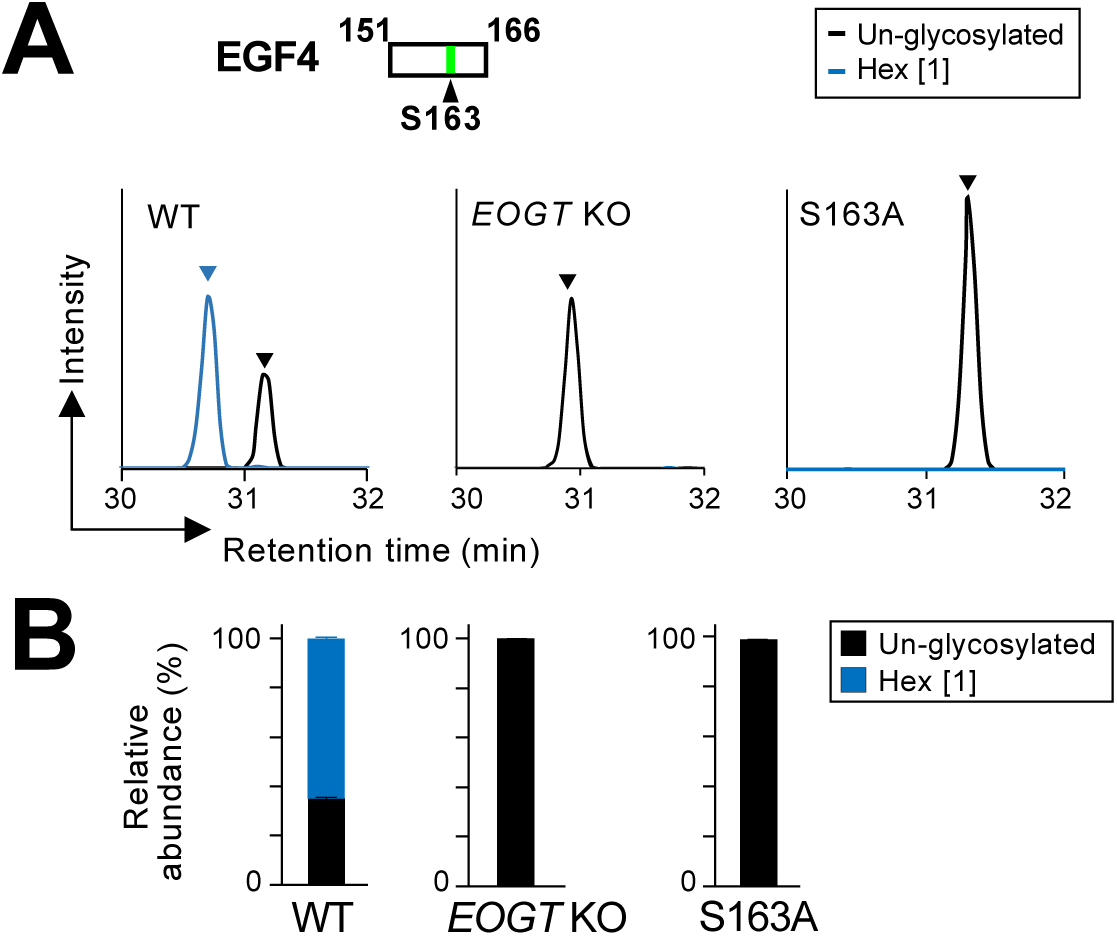
O-hexose on EGF4 is modified by EOGT. **(A)** Protease digests of DLK1 ECD isolated from the culture medium of HEK293T cells (WT) or their *EOGT*-deficient cells (*EOGT* KO) were analyzed by LC-MS/MS. The DLK1 ECD S163A mutant produced in HEK293T cells were also subjected to analysis. The extracted ion chromatograms reveal the presence or absence of an O-Hex glycoform modifying EGF4 of DLK1 ECD. Arrowheads indicate peaks of assigned (glyco)peptide spectra. The MS/MS spectra are presented in Fig. S6. **(B)** Semiquantitative analysis of respective glycoforms presented in (A). The relative abundance of respective glycoforms was calculated from the integrated peak height values and are presented as the mean ± the range of data (n=2).

### Generation of HEK293 cell lines lacking POFUT1, POGLUT1, or EOGT

To gain insight into the roles of O-glycans that modify DLK1, we generated cell lines lacking EGF domain-specific glycosyltransferases using CRISPR/Cas9-mediated genome editing. Although HEK293T cell lines lacking *POFUT1* (34), *POGLUT1* (34), or *EOGT* (44) have been generated previously, we chose to use HEK293 cells because they adhere loosely to culture plates and are suitable for downstream flow cytometry experiments. A deleted genomic region was identified (Figure S22). The protein expression of POFUT1 and EOGT was abolished in *POFUT1*- and *EOGT*-deficient cells (Figure 4A). The *POGLUT1*-deficient cells were confirmed by the loss of mRNA expression of POGLUT1 (Figure 4B). *POFUT1*-, *POGLUT1*-, and *EOGT*-deficient cells derived from HEK293 cells were used for further experiments.

**Figure 4.**
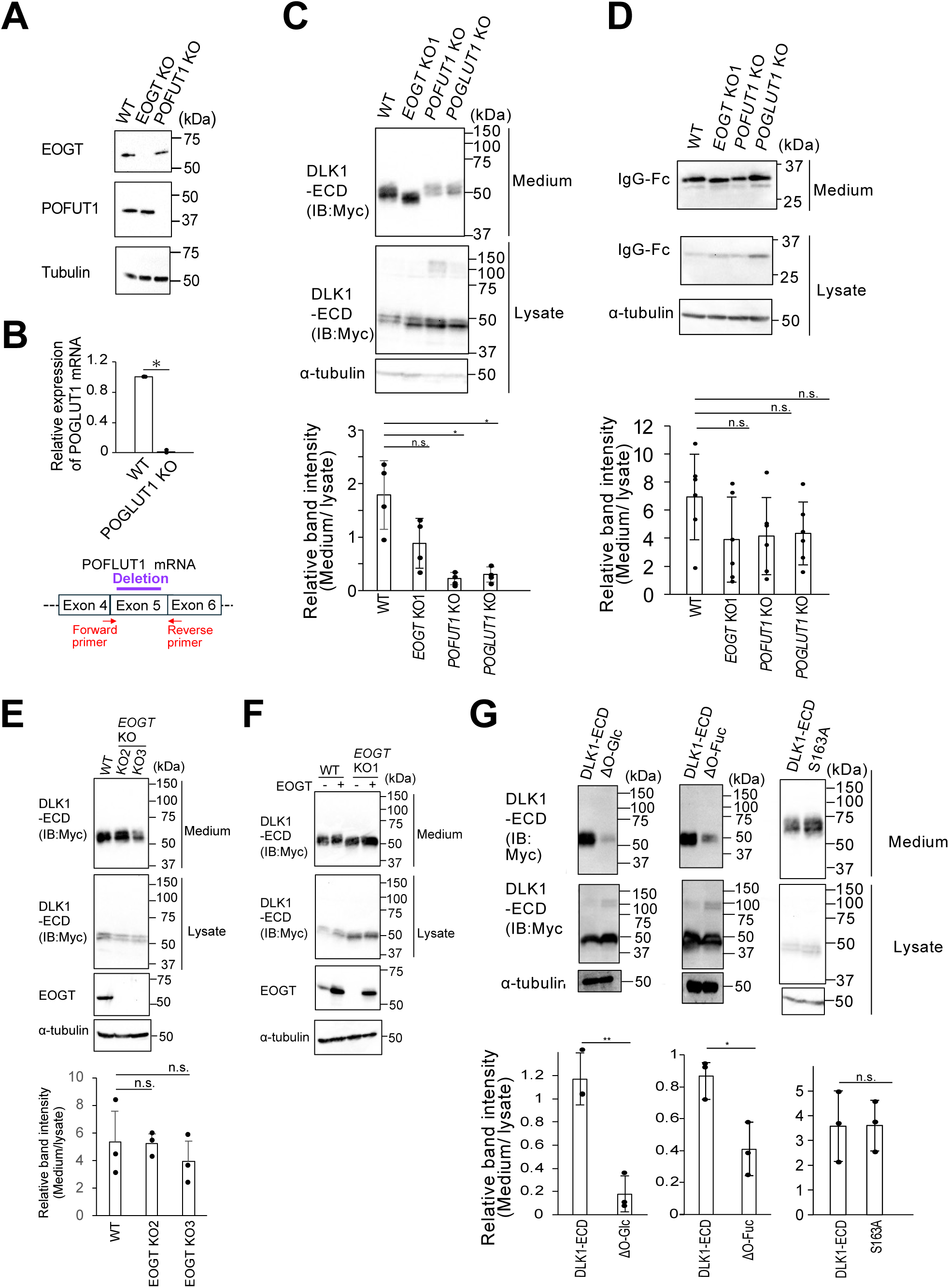
Lack of O-Glc or O-Fuc impairs the secretion of DLK1 ECD. **(A)** Expression of EOGT and POFUT1 in *EOGT*-deficient cells (*EOGT* KO1), *POFUT1*-deficient cells (*POFUT1* KO) derived from HEK293 cells. The cell lysate was separated by SDS-PAGE and detected by each antibody indicated. **(B)** Relative mRNA expression of *POGLUT1* in *POGLUT1* KO cells and wild-type HEK293 cells. Values represent the mean ± standard deviation (s.d.), n=3. Statistical analysis was performed using one-way ANOVA and Tukey’s multiple-comparison test. \**p*<0.05. n.s., not significant. **(C)** DLK1 ECD was expressed in wild-type HEK293 cells (WT), *EOGT*-deficient cells (*EOGT* KO1), *POFUT1*-deficient cells (*POFUT1* KO), or *POGLUT1*-deficient cells (*POGLUT1* KO). The expression of DLK1 ECD in the culture medium and the cell lysate was evaluated by immunoblotting with indicated antibodies. Presented below is quantitative analysis of DLK1 ECD secretion. The band intensity of secreted DLK1 ECD was divided by that from cell lysates. Values represent the mean ± s.d., n=4. Statistical analysis was performed using one-way ANOVA and Tukey’s multiple-comparison test. \**p*<0.05. n.s., not significant. **(D)** Same as (C) but for human IgG-Fc. Values represent the mean ± s.d., n=6. **(E)** Secretion of DLK1 ECD from *EOGT* KO2 and KO3 clones. The expression of DLK1 ECD in culture medium and cell lysate was evaluated as indicated above. A statistically significant difference was estimated by Student’ s *t*-test. n.s., not significant **(F)** Restoring EOGT expression in *EOGT* KO1 cells. *EOGT* KO1 or HEK293 cells were transfected to express DLK1 ECD in the presence or absence of exogenously expressed EOGT. **(G)** HEK293 cells were transfected to express DLK1 ECD and its variants harboring S94A/S214A mutations at O-glucosylation sites (ΔGlc), T143V/T222V mutations at O-fucosylation sites (ΔFuc), or S163A mutation. Values represent mean ± s.d., *n*=3. A statistically significant difference was estimated by Student’ s *t*-test. n.s, not significant; \**p*<0.05; \*\**p*<0.01.

### O-fucosylation and O-glucosylation promote secretion of DLK1-ECD

As immature proteins tend to be retained in the ER, we expressed a soluble DLK1 construct in HEK293 cells and evaluated its secretion efficiency by calculating the ratio of DLK1 in the culture medium to that in the cell lysate. As presented in Figure 4C, DLK1-ECD was secreted less efficiently in *POFUT1* or *POGLUT1* mutant cells than in control HEK293 cells. In contrast, the secretion efficiency of DLK1-ECD from *EOGT*-deficient cells was comparable to that of control cells. However, the absence of POFUT1, POGLUT1, or EOGT did not affect the secretion of IgG-Fc lacking the EGF domains (Figure 4D). Although an increase in the mobility of DLK1-ECD was observed in *EOG*T-KO1 cells, independent *EOGT* KO clones (KO2 and KO3) did not exhibit a mobility shift in DLK1-ECD compared to wild-type HEK293 cells (Figure 4E). Furthermore, the restoration of EOGT expression in *EOG*T-KO1 cells did not change the mobility of DLK1-ECD (Figure 4F). Therefore, the change in the migration of DLK1-ECD was specific to a particular clone. Nonetheless, these results consistently indicated that EOGT is dispensable for DLK1-ECD secretion.

To pinpoint the action of O-glycosyltransferase activity toward DLK1, we generated the ΔGlc and ΔFuc mutant variants for DLK1-ECD, in which the two major modification sites for POGLUT1 (i.e., Ser94 and Ser214) or POFUT1 (i.e., Thr143 and Thr222) were substituted by changing Ser to Ala or Thr to Val. As presented in Figure 4G, the ΔGlc and ΔFuc mutants exhibit poor secretion efficiency compared to that of control DLK1-ECD. In contrast, the S163A mutant lacking EOGT-dependent O-Hex modification secreted at a level comparable to that of the control. These data suggested that POGLUT1 and POFUT1 play pivotal roles in the effective secretion of DLK1-ECD.

### Establishment of a transport assay for DLK1 trafficking from the ER to the cell surface

The decreased secretion of DLK1-ECD may be due to a delay in the folding process and/or subsequent trafficking from the ER to the cell surface. To monitor the DLK1 transport from the ER to the cell surface, we used a modified version of the retention system with a selective hook (RUSH) system (48). In this system, the green fluorescent protein (GFP) and streptavidin-binding peptide (SBP) were fused to the C-terminus of FLAG-DLK1 (Figure 1D). As a binding partner (Hook protein), streptavidin (Str) and HA-tag were fused with a fragment of the H-2 class II histocompatibility antigen gamma chain known as Ii (CD74) that is targeted to the ER (Figure 5A). We confirmed that doxycycline treatment induced concomitant Reporter DLK1 and Hook protein expression (Figure 5B-D). Both proteins exhibited perinuclear ER-like localization and physically interacted with each other (data not indicated). Upon biotin supplementation, the Reporter DLK1 emerged on the cell surface, and its expression reached its highest level at 40 min (Figure 5C).

**Figure 5.**
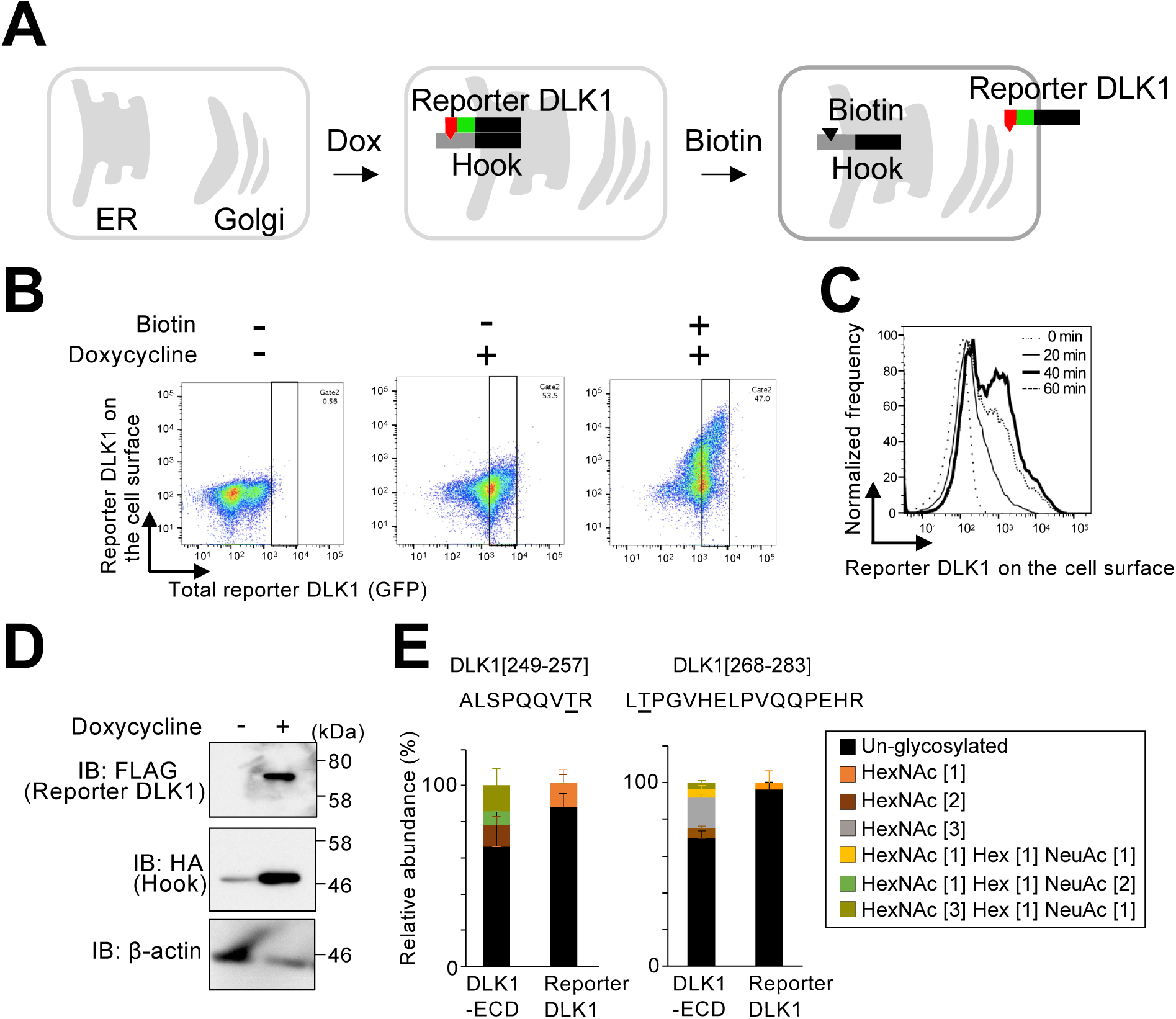
Establishment of a cell surface transport assay for DLK1. **(A)** The cell surface transport assay for DLK1 was developed based on the RUSH method. Reporter DLK1 is composed of DLK1 (*black*), GFP (*green*), and streptavidin-binding peptide (SBP, *red*), while Hook protein is composed of Ii (*black*) and streptavidin (*grey*). Reporter DLK1 and Hook protein were designed so that both proteins bind via SBP and streptavidin fused to the C-terminus of Reporter DLK1 and Hook protein, respectively. Upon induction with doxycycline (DOX), the Ii domain directs Hook protein to the ER where Reporter DLK1 is accumulated. Biotin competes for Hook protein, thus allowing Reporter DLK1 to be transported to the cell surface via the Golgi. **(B)** Expression of Reporter DLK1 in the transport assay. After DOX induction, transfected HEK293 cells were treated with biotin for 60 min and analyzed by flow cytometer. Anti-DLK1 antibody detects cell surface expression of Reporter DLK1. GFP expression reflects the total expression level of Reporter DLK1. **(C)** Time course for the surface expression of Reporter DLK1 after the addition of biotin. Transfected HEK293 cells were gated at a constant GFP intensity in all samples. **(D)** Detection of Reporter DLK1^del^ and Hook protein expressed in HEK293 cells. **(E)** Comparison of mucin type O-glycan modifications on DLK1^[249-257]^ and DLK1^[268-283]^ peptides derived from the DLK1-ECD or Reporter DLK1. The mean ± standard deviation (n=3) is presented. A list of glycopeptides detected by mass spectrometry is summarized in Table S1 (DLK1-ECD) and Table S2 (Reporter DLK1). The MS/MS spectra are presented in Fig.S7 and S8 (DLK1-ECD) and Fig.S9 and S10 (Reporter DLK1).

The retention of the Reporter DLK1 by Hook protein was confirmed independently by detecting mucin-type O-glycans. This is based on the knowledge that O-N-acetyl-galactosamine (O-GalNAc) modification and elongation typically occur in the Golgi. LC-MS/MS analysis of the Reporter DLK1 co-expressed with the Hook protein revealed minimal O-GalNAc modifications, and elongated structures were barely detectable at Thr256 and Thr269 (Figure 5E). This contrasts with the secreted DLK1-ECD, supporting the ER distribution of the Reporter DLK1.

Considering the potential processing of Reporter DLK1 on the cell surface, the reporter construct was further modified by deleting the juxtamembrane region (Glu279 - Glu301) that is cleaved during ectodomain shedding by tumor necrosis factor-α converting enzyme (TACE/ADAM17) (49). The deletion mutant (Reporter DLK1^del^) was similarly detected on the cell surface (Figure 6A) and its expression increased in a time-dependent manner until 60 min. Based on this observation, we used the Reporter DLK1^del^ for further analysis.

**Figure 6.**
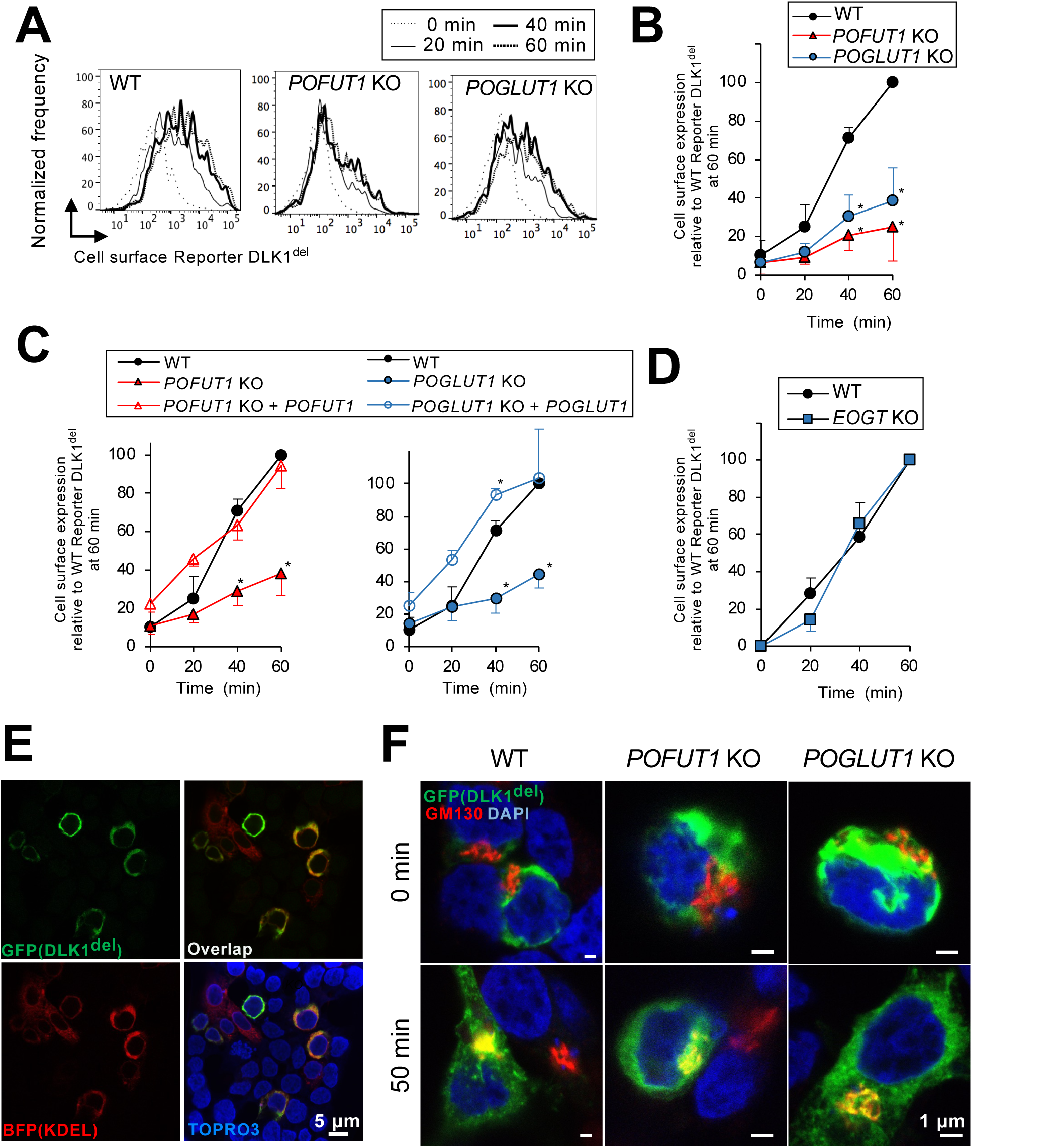
Cell surface transport of Reporter DLK1 from the ER is perturbed in *POFUT1*- or *POGLUT1*-deficient HEK293 cells. **(A)** Cell surface transport assay for Reporter DLK1^del^ expressed in HEK293 cells (WT), *POFUT*1-deficinet cells (*POFUT1* KO), or *POGLUT*1-deficinet cells (*POGLUT1* KO). After induction with doxycycline, cells were incubated with biotin for the indicated times and then stained for DLK1 for analysis using a flow cytometer. The expression levels of Reporter DLK1^del^ in all samples were matched by gating GFP-positive cells at a constant intensity. **(B)** Comparison of the surface expression of Reporter DLK1^del^ on the cell surface of HEK293 cells, *POFUT*1-deficinet cells, and *POGLUT*1-deficinet cells. The intensity of the signals obtained by anti-DLK1 antibody was normalized against that of HEK293 cells at 60 min. Values represent the mean standard deviation (s.d.), *n*=3. Statistical analysis was performed using two-way ANOVA and Tukey’s multiple-comparison test, \**p*<0.05 (WT versus *POFUT1* KO or *POGLUT1* KO). **(C)** Same as in (B) but with the rescued cells with POFUT1 or POGLUT1. Values represent the mean ± s.d., *n*=3. The statistically significant difference against WT was estimated by two-way ANOVA and Tukey’s multiple-comparison test, \**p*<0.05. **(D)** Same as in (B) but for *EOGT*-deficient cells (*EOGT* KO). Values represent the mean ± s.d., *n*=3. **(E)** Subcellular localization of Reporter DLK1^del^. Reporter DLK1^del^ (*green*), Hook protein, and BFP-KDEL (*red*) were co-expressed in HEK293 cells. Cells treated with doxycycline were fixed and mounted in TOPRO3-containing media to stain the nucleus (*blue*). Note that DLK1^del^ was retained in the ER in the absence of biotin. **(F)** Change in the subcellular localization of Reporter DLK1^del^ (*green*) after addition of biotin. Doxycycline-treated cells were incubated with biotin for the indicated time. Fixed cells were then stained with an anti-GM130 antibody to label the Golgi (*red*), and DAPI was used to label the nucleus (*blue*).

### O-fucose and O-glucose glycans on DLK1 are important for the proper transport of DLK1 from the ER to the cell surface

To determine if POFUT1 or POGLUT1 affected the trafficking of the Reporter DLK1^del^ from the ER to the cell surface, a transport assay was performed using *POGLUT1*-or *POFUT1*-deficint cells (Figure 6A-D). For a proper comparison of different cell lines, the expression levels of the Reporter DLK1^del^ in all samples were matched by gating the GFP-positive cells at a constant intensity. Flow cytometry revealed that both mutants exhibited decreased cell surface expression of the Reporter DLK1^del^. The rescue experiment further supported that POFUT1 and POGLUT1 are responsible for promoting the trafficking of the Reporter DLK1^del^ from the ER to the cell surface. In parallel with the secretion assay, the Reporter DLK1^del^ transport was unaffected by the lack of EOGT (Figure 6D).

To visualize the subcellular localization of the Reporter DLK1^del^ during the transport assay, co-localization of the Reporter DLK1^del^ with the ER marker BFP-KDEL or Golgi marker GM130 was analyzed. Prior to biotin supplementation, the Reporter DLK1^del^ largely co-localized with BFP-KDEL in wild-type HEK293 cells (Figure 6E). In contrast, after the addition of biotin for 50 min, Golgi staining, marked by GM130, relative to perinuclear ER-like staining, was more prominent in wild-type HEK293 cells than in *POGLUT1*- or *POFUT1*-deficient cells (Figure 6F). These data suggest that impaired transport from the ER, rather than from the Golgi, accounts for decreased cell surface transport in *POGLUT1*- or *POFUT1*-deficient cells.

### O-glycan analysis of Reporter DLK1 tethered to the ER

As EGF domains can be modified by multiple types of glycans, the absence of POGLUT1 or POFUT1 could affect the glycosylation level of the remaining glycans. To assess the glycosylation status of ER-bound DLK1, the reporter DLK1 purified from wild-type HEK293 cells was compared to that purified from *POGLUT1* or *POFUT1* mutant HEK293 cells (Table 2 and Figure 7A). Overall, the glycosylation pattern of the Reporter DLK1 from wild-type HEK293 cells was similar to that of DLK-ECD, except for a higher level of N-glycosylation at EGF4, presumably due to extended ER retention and post-translational modification of skipped N-glycosylation sequences (50, 51).

**Table 2.**
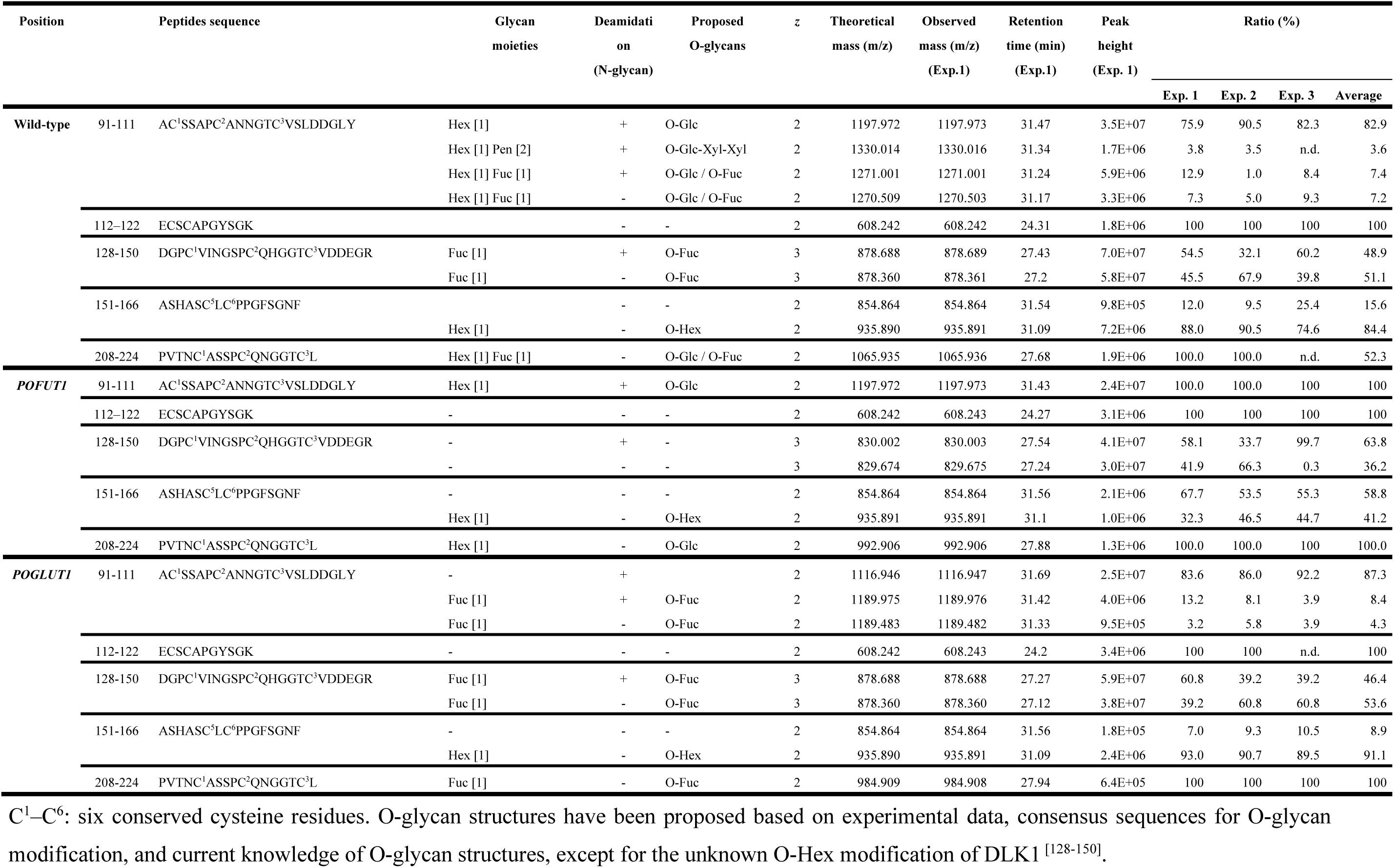
List of proteolytic fragments of Reporter DLK1 containing putative glycosylation sites for EGF domain-specific O-glycosylation.

**Figure 7.**
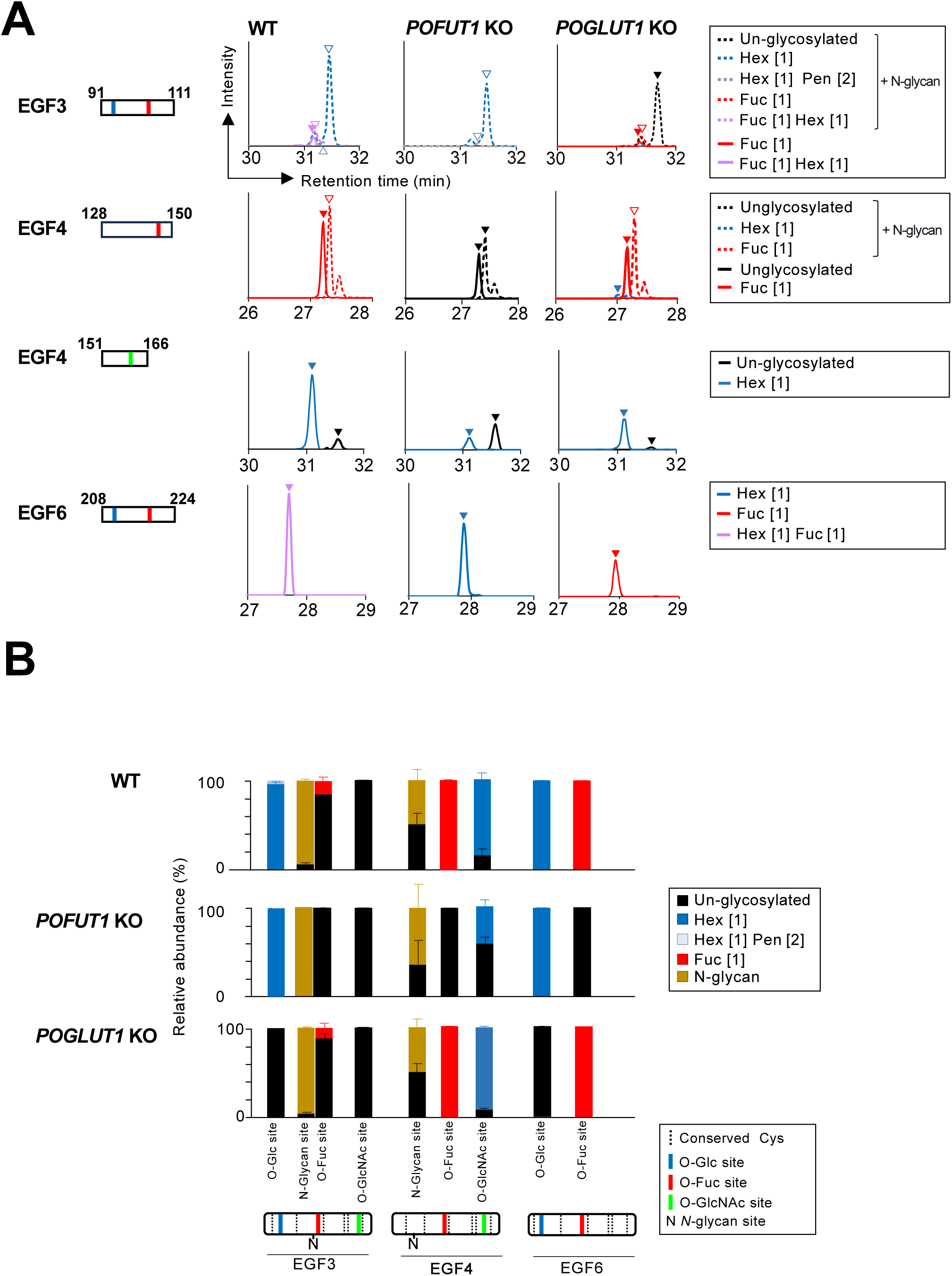
O-glycosylation profile of ER-tethered Reporter DLK1 expressed in *POFUT1* or *POGLUT1*-deficinet HEK293 cells. **(A)** LC-MS/MS analysis of protease digests of Reporter DLK1 expressed in wild-type HEK293 cells (WT), the *POFUT1*-deficient cells (*POFUT1* KO), or the *POGLUT1*-deficient cells (*POGLUT1* KO). Reporter DLK1 was retained in the ER by co-expressing with Hook protein. Extracted ion chromatograms (EICs) of unglycosylated peptides (*black*) and glycopeptides carrying O-glycans (indicated by different colors) are presented. Arrowheads indicate the position of the precursor ions in the annotated MS/MS spectra (Fig.S11-21). The glycopeptides detected by mass spectrometry are summarized in Table 2. **(B)** Relative abundance of O-glycans detected on Reporter DLK1 produced in wild-type HEK293, *POFUT1*-deficient cells, or *POGLUT1*-deficient cells. The relative abundance of respective glycoforms was calculated from the integrated peak height values and is presented as the mean ± standard deviation, n=3. The DLK1 domain structure is indicated below with predicted glycan modification sites: O-Fuc (*red*), O-Glc (*blue*), and O-GlcNAc (*green*).

When the Reporter DLK1 was expressed in *POFUT1*-deficient cells, dHex (i.e., Fuc) modifications of EGF3, EGF4, and EGF6 were abolished, as expected from the substrate specificity of POFUT1. Similarly, Hex modifications assigned to the O-Glc modification sites were undetectable in EGF3 and EGF6 in *POGLUT1*-mutant cells (Figure 7B). Importantly, the lack of O-Fuc did not affect the state of O-Glc modification, and vice versa. Given that the activities of POFUT1 and POGLUT1 depend on the folded state of the EGF domains, these findings suggest that the Reporter DLK1^del^ complete the folding process required for O-glycosylation in the ER before being transported to the cell surface.

Notably, although the lack of O-glycan did not affect the overall level of DLK1 N-glycosylation, the small population of EGF3 unmodified by N-glycan was diminished in *POFUT1*-deficient cells, as expected from the mutually exclusive relationship between N-glycosylation and O-fucosylation on EGF3. It is also noted that the EOGT-dependent O-Hex modification of EGF4 is slightly but significantly reduced in *POFUT1*-deficient cells (Figure 7B) (one-way ANOVA, Tukey’s Multiple Comparison, p < 0.01).

### Decreased turnover of DLK1 in the absence of O-glycosyltransferases

To analyze the effects of the lack of O-Fuc or O-Glc glycans on DLK1 expression, FLAG-tagged full-length DLK1 (FLAG-DLK1) was transiently expressed in wild-type and mutant HEK293 cells, and a chase experiment was performed by inhibiting new protein synthesis with cycloheximide. In wild-type cells, the FLAG-DLK1 levels decreased rapidly within 1 h (Figure 8A). In contrast, the relative lifetime of FLAG-DLK1 was significantly prolonged in the mutants lacking *POFUT1* or *POGLUT1* (Figure 8EB). These results suggest that O-glycans are critical for DLK1 proteostasis, and this is in agreement with the transport and secretion assays.

**Figure 8.**
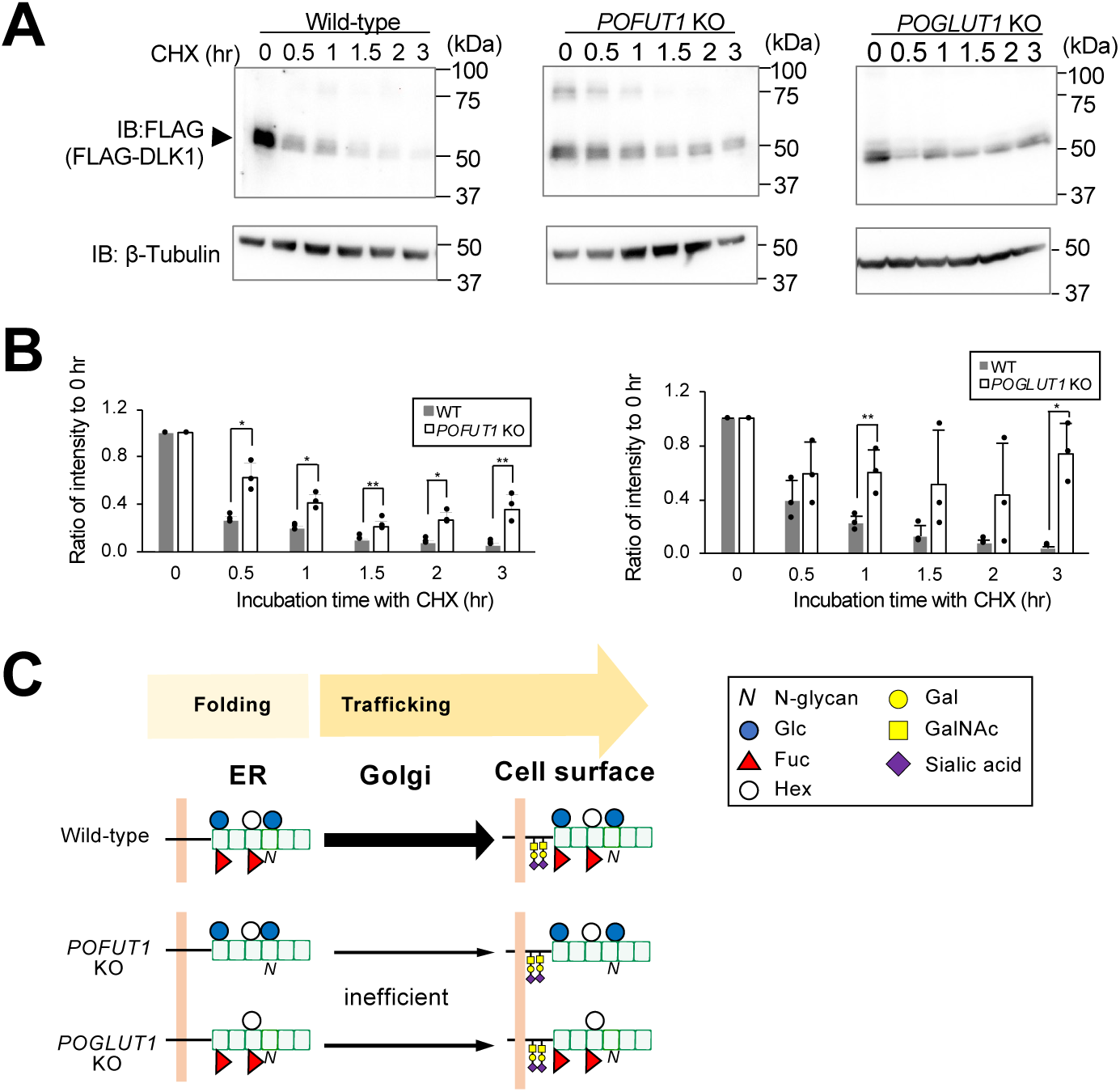
Effect of respective O-glycosyltransferase on the turnover of DLK1. **(A)** Time course of FLAG-DLK1 transiently transfected into wild-type HEK293 cells (WT) compared to that transfected into *POFUT1*-deficient cells (*POFUT1* KO), or *POGLUT1*-deficient cells (*POGLUT1* KO). Cells were incubated with cycloheximide (CHX) for the indicated time. The whole cell lysate was prepared and subjected to SDS-PAGE and immunoblotting with anti-FLAG tag or anti-β-tubulin antibody. **(B)** Quantification of the relative DLK1 band intensity upon cycloheximide treatment. The band intensity of FLAG-DLK1 was normalized using that of β-tubulin as a reference. Values are expressed as the mean ± standard deviation (s.d.), n=3. Statistical significance was assessed using Student’ s *t*-test. \**p*<0.05, \*\**p*<0.01. **(C)** Model for the roles of O-glycan during DLK1 trafficking to the cell surface. DLK1 is modified with O-glycans, including O-Fuc, O-Glc, and newly identified O-Hex, through the action of POFUT1, POGLUT1, and EOGT, respectively. Lack of *POFUT1* or *POGLUT1* did not result in an apparent loss of remaining O-glycans, indicating the completion of the folding processes. Transport assay revealed that lack of POFUT1 or POGLUT1 but not EOGT perturbed the transport of DLK1 from the ER. O-GalNAc modification occurs after exit from the ER.

## Discussion

The critical role of O-glycans in EGF domains during protein maturation has been widely recognized in the study of Notch receptors. The lack of O-glycans results in ER retention of Notch receptors (52, 53). Similarly, O-glycans are important for secretion of Notch ectodomains. As a molecular mechanism, it has been proposed that O-glycans stabilize EGF domains and thus promote EGF domain folding (34–38). The effect of O-glycans on protein secretion and stabilization was further supported by studies examining thrombospondin type 1 repeat (TSR) domain-containing proteins such as TSP1 and ADAMTSL1 (54). However, O-glycans may promote trafficking from the ER independent of their role in stabilization. In this study, we directly investigated whether O-glycans affect trafficking using ER-tethered DLK1 in the RUSH system.

Our comprehensive mass spectrometric analysis provided information on the folding state of DLK1, ultimately indicating no evidence of misfolding of the ER-tethered DLK1 in the absence of POFUT1 or POGLUT1. Therefore, decreased cell surface expression of the Reporter DLK1 in *POFUT1*- or *POGLUT1*-deficient cells is likely due to defects in DLK1 transport rather than impaired protein folding. The molecular mechanisms by which O-glycans affect DLK1 trafficking via secretory pathways remain unknown and multiple mechanisms involving anterograde or retrograde protein transport should be considered. Nonetheless, our results provide the first evidence that O-glycans affect protein trafficking from the ER independently of the folding process required for O-glycosylation in the ER (Figure 8F).

Semi-quantitative site-specific glycoproteomics of DLK1-ECD has revised the current view of the substrate specificity of EOGT. A previous study examining multiple O-GlcNAc-modified EGF domains of NOTCH1 revealed that C^5^-X-X-G/S/P-Y/F/T-T/S-G-X-X-C^6^ was the preferred sequence for O-GlcNAcylation at higher stoichiometry (32). However, this study revealed that the two sites in DLK1, EGF3, and EGF4 that satisfied the aforementioned sequence failed to be modified with O-GlcNAc. Therefore, the number of potentially O-GlcNAcylated EGF domains was smaller than expected. Furthermore, we identified a novel O-Hex modification at the O-GlcNAc modification site in DLK1 EGF4. DLK1 represents the first example of an O-Hex modification of O-GlcNAc sites, although there is an implication that trace amounts of O-Hex may be present at the O-GlcNAc sites of NOTCH3 (36). Although the identity of the Hex modification has not been directly addressed, we detected weak O-glucose transferase activity toward chemically synthesized *Drosophila* EGF20 (dEGF20) *in vitro* (33) (Figure S23). Whether EOGT catalyzes the O-Glc modification requires further investigation. Notably, O-Hex modification was slightly decreased in the Reporter DLK1 produced by *POFUT1* mutant cells. Given that no physical interactions have been reported between different O-glycosyltransferases, we speculated that O-fucosylation might induce conformational changes in DLK1 EGF4 that favor O-Hex modification by EOGT.

Another new insight gained from this study was the potential interplay between N-glycosylation and O-fucosylation. The amino acid sequence of DLK1 EGF3 contains a consensus sequence for O-fucosylation (NGT̅C^3^) but overlaps with the sequence for N-glycosylation (NXT) (Figure 2C). In DLK1-ECD secreted into the culture medium, N-glycosylation and O-fucosylation of EGF3 were nearly mutually exclusive. Based on the current view that N-glycosylation can occur on nascent proteins (50, 51) but that O-fucosylation occurs after the folding of EGF domains, it is likely that N-glycan in the vicinity of the O-fucosylation site inhibits the enzymatic activity of POFUT1, possibly through steric hindrance.

While extensive research has been conducted to examine NOTCH receptors, the roles of O-glycans in other structurally related proteins, such as Notch ligands, have only been investigated in a few studies (55, 56). Delta-like protein 1 (DLL1), a canonical ligand of NOTCH signaling, is O-fucosylated at multiple sites, specifically in four of the eight EGF domains (56). In cells lacking *POFUT1*, DLL1 accumulated intracellularly when expressed exogenously or endogenously. However, the exact subcellular localization of DLL1 has not yet been experimentally determined, although its cell surface expression appears normal (56). As suggested in this study, the inefficient transport of DLL1 from the ER to the cell surface may contribute to its abnormal intracellular accumulation.

Unlike Notch receptors, O-glycans on DLK1 isolated from HEK293T cells are predominantly monosaccharide, which is consistent with the results for mouse DLK1 isolated from amniotic fluid. However, the extension of O-glycans is partly dependent on the expression of glycosyltransferases that mediate O-glycan elongation. Further investigation is needed to determine whether O-glycan extension occurs on the EGF domains of DLK1 under different cellular conditions. Regardless, the O-glycosylation map of DLK1 EGF domains revealed in this study serves as a blueprint for analyzing changes in EGF domain-specific O-glycans and their associated molecular functions during DLK1-dependent biological processes.

## Experimental Procedures

### Antibodies

The antibodies used for this study included anti-human DLK1 antibody clone 211309 (R&D Systems Minneapolis, MN, USA), anti-DYKDDDDK-tag antibody clone 1E6 (FUJIFILM Wako Pure Chemical Corporation, Osaka, Japan), anti-DYKDDDDK-tag antibody clone 15 (Biolegend), anti-EOGT (AER61) antibody (Abcam, Cambridge, UK), Anti-POFUT1 (Proteintech, IL, USA), anti-GM130 antibody (Becton, Dickinson and Company (31), NJ, USA), anti-HA antibody clone 4B2 (FUJIFILM Wako Pure Chemical Corporation), anti-HA antibody clone C29F4 (CST, MA, USA), anti-Myc antibody clone 9E10 (DSHB), anti-α-tubulin antibody clone 12G10 (DSHB, IA, USA), PE-conjugated goat anti-mouse IgG (#405307), horseradish peroxidase (HRP)-conjugated anti-mouse IgG, HRP-conjugated anti-rabbit IgG antibody (CST; #7074P2), HRP-conjugated anti-human IgG-Fc (Bethyl Laboratories, Inc, TX, USA), HRP-conjugated anti-rat IgG antibody (#SA00001-15; Thermo Fisher Scientific, MA, USA), and Alexa 594-conjugated anti-mouse IgG (Invitrogen). The antibody dilutions used in the experiments are summarized in Table S4.

### Plasmids

The Str-Ii_SBP-mCherry-Golgin84 plasmid was gifted by Franck Perez (Addgene Plasmid # 65304) (57). The BFP-KDEL plasmid was gifted by Gia Voeltz (Addgene Plasmid # 49150) (58). The pCX-bsr-human_ManII-G4S2×8-EGFP-SBP and pCMV-hyPBase-teton-P2A-BSD vectors were a gift from Yusuke Maeda of Osaka University, Japan. The original pCMV-hyPBase vector was provided by the Trust Sanger Institute (UK) (59). Expression vectors for mouse Pofut1:MycDDK (#MR206140) and mouse Poglut1:MycDDK (#MR206123) were purchased from OriGene (MD, USA). The expression vector for human IgG-Fc has been described previously (34). The expression vector for mouse EOGT-IRES-GFP has been previously used (60).

### Plasmid constructs

The human DLK1 gene used in this study is a natural variant (VAR_060335) with an R101G sequence (22, 61). To generate a vector expressing the human DLK1 ectodomain (ECD), the entire EGF domain and the flanking region modified by mucin-type O-glycans (amino acid position 18-302) were inserted into the pSeqTag2C-MycHis6 vector (Invitrogen) at the SfiI/NotI sites. Site-specific mutagenesis of DLK1 was performed using PCR according to the protocol of the PrimeSTAR mutagenesis basal kit (Takara, Shiga, Japan). The primer sequences are listed in Table S5.

To generate a vector expressing FLAG-DLK1, the EcoRI/AfeI fragment of a synthetic oligonucleotide containing the Igκ signal sequences and FLAG tag sequence and the AfeI/XbaI fragment of human DLK1 (amino acid sequence 24-383) were inserted into the EcoRI/XbaI sites of the pTracer-CMV vector. The resultant pTracer-CMV-FLAG-DLK1 vector served as a template for preparing two PCR fragments designed to exclude the Glu279-Glu301 region. The AfeI/ClaI and ClaI/XbaI fragments were simultaneously ligated to the AfeI/XbaI sites of the pTracer CMV vector to generate the FLAG-DLK1^del^ construct.

To construct the IRES-containing bicistronic vector encoding Ii fused to streptavidin (Str) (thereafter called as Hook protein) and DLK1 fused to EGFP and streptavidin-binding peptide (SBP) (thereafter called as Reporter DLK1), pCMV-hyPBase-teton-P2A-BSD vector was inserted into EcoRI/HpaI sites with the following three fragments: i) EcoRI/NheI digests of Str-Ii-IRES fragments amplified from Str-Ii_SBP-mCherry-Golgin84 vector; ii) NheI digests of FLAG-DLK1 amplified from pTracer-CMV-FLAG-DLK1 vector; iii) NheI/HpaI digests of G4S2×8-EGFP-SBP fragments amplified from pCX-bsr-human_ ManII-G4S2×8-EGFP-SBP vector. Reporter DLK1^del^ construct was generated by swapping an NheI fragment encoding FLAG-DLK1. The primer sequences are listed in Table S6.

### Cell lines

Human embryonic kidney (HEK) 293 cells were obtained from the RIKEN BioResource Research Center. *EOGT*-deficient HEK293T cells were generated as described previously (44). The cells were cultured in Dulbecco’s modified Eagle’s medium (DMEM) supplemented with heat-inactivated fetal bovine serum (FBS) and penicillin-streptomycin.

To generate HEK293 cells stably expressing DLK1-ECD, cells were transfected with pSecTag-DLK1-ECD and selected with 800 μg/mL hygromycin B (Fujifilm). To generate HEK293 and its derivative cells expressing Reporter DLK1 and Hook protein, selection was performed in the presence of 8 μg/mL blasticidin S hydrochloride (FUJIFILM Wako Pure Chemical Corporation).

*POFUT1-*, *POGLUT1-*, and *EOGT-*deficient HEK293 cells were generated using the CRISPR-Cas9 gene editing system. pX330-CAS9-mGFP-sgRNA constructs were generated from the pX330-CAS9-mGFP (59) plasmid by inserting a specific sgRNA target sequence, as described in Table S7. The resulting plasmids were transfected into HEK293 cells using polyethyleneimine (PEI)-Max MW 40,000 (Polysciences). After 1.5 days, GFP-positive cells were sorted using a FACS Aria Fusion cell sorter (Becton, Dickinson and Company). The sorted cells were subjected to limited dilution and individual clones were selected and analyzed for their genomic sequences.

### Quantitative real-time polymerase chain reaction (qPCR)

Total RNA was extracted using QIAzol Lysis Reagent (QIAGEN, Hilden, Germany) and subjected to reverse transcription using the Prime Script RT reagent kit (Takara). The generated cDNA was analyzed by qPCR using the SYBR Green qPCR Master Mix (Bio-Rad), and the primer sets are listed in Table S8.

### Transfection

The cells were incubated in a fresh culture medium without penicillin-streptomycin. After 2 h, the cells were transfected with PEI-Max as previously described (36). Briefly, PEI-Max and plasmid DNA were individually diluted in Opti-MEM (Gibco/Thermo Fisher Scientific) and combined to form a PEI-Max/plasmid DNA complex. After a 15-min incubation at room temperature (RT), the mixtures were added to the cells. After one day, the medium was replaced with a complete cell culture medium.

### Purification of DLK1-ECD and Reporter DLK1

To purify DLK1-ECD, HEK293T cells were transiently transfected into 10 cm or 15 cm dishes using PEI-max. Transfected cells were cultured in a complete cell culture medium for four days. After collecting the spent medium, the cells were cultured in complete medium for an additional three days. The spent medium was centrifuged at 400 *g* and 4°C for 4 min. The supernatant was incubated with 40 μL (bed volume) of anti-His-tag mAb-magnetic beads (MBL) in a cold room overnight. After centrifugation at 400 *g* at 4°C for 1 min, the supernatant was removed, and the beads were washed five times with 0.5 mL of a solution containing 50 mM Tris-HCl (pH 7.4), 150 mM NaCl, and 0.1% NP-40. The bound protein on the beads were eluted by heating the samples at 95°C for 5 min in 90 μL of SDS sample buffer (133 mM Tris-HCl, pH6.8, 2.2% SDS, 11.1% glycerol, and 0.01% bromophenol blue).

DLK1 was purified from cells transfected with plasmids encoding the hook protein and Reporter DLK1. Briefly, cells grown on ten 15 cm dishes were treated with 1 μg/μL doxycycline in the complete culture medium. After protein induction for 1.5 days, the cells were lysed in 15 mL of lysis buffer (20 mM Tris-HCl, pH 7.4, 150 mM NaCl, and 1% Triton X-100) with protease inhibitors. The lysate was incubated on ice for 30 min and then centrifuged at 18,000 *g* at 4°C for 15 min. The supernatant was incubated with 50 μL (bed volume) of DDDDK-tagged protein purification gel (MoAb clone FLA, MBL) in a cold room overnight. The gels were washed five times with a solution containing 50 mM Tris-HCl (pH 7.4), 150 mM NaCl, and 0.1% NP-40 followed by elution with 30 μL of 2x SDS sample buffer as described above.

### Sample preparation for mass spectrometry

The samples were reduced with 90 mM tris(2-carboxyethyl)phosphine (Thermo Fisher Scientific) at 100°C for 5 min. For alkylation, the samples were incubated with 360 mM iodoacetamide dissolved in 50 mM Tris-HCl (pH 6.8) for 1 h at RT in the dark. The samples were subjected to SDS-PAGE and stained with GelCode Blue Stain Reagent (Thermo Fisher Scientific). Stained bands were excised and transferred to low-binding protein tubes (InaOptica, Osaka, Japan). The gels were washed with 1 mL of 50% methanol and 20 mM diammonium phosphate (DiAP) for 1h at RT with rotation. Subsequently, the gels were washed again with 50% methanol and 20 mM DiAP and rotated at 4°C overnight. The washing step was repeated thrice with 50% methanol and 20 mM DiAP for 30 min at RT with rotation. Acetonitrile was added to the gel, incubated for 30 min, and dried.

Samples were incubated with 30 ng/µL trypsin (Promega) diluted in 20 mM DiAP for 5h at 37°C. The supernatant was collected, and the digested peptides were added to 50 µL Milli-Q water and incubated at RT for 10 min. They were then supplemented with 33 µL of acetonitrile containing 0.1% formic acid and incubated for 20 min at RT. The samples were further treated with 42 µL of acetonitrile containing 0.1% formic acid and incubated for 20 min at RT. The supernatant was collected and subjected to vacuum freeze-drying to reduce its volume. The samples were then added to ice-cold acetone (four times the sample volume). After incubating at −30°C overnight, the samples were centrifuged at 18,000 *g* for 15 min at 4°C. The supernatant was removed and the pellet was dried using a vacuum freeze dryer.

Tryptic fragments were further digested with chymotrypsin by dissolving the samples in 20 mM ammonium carbohydrate containing 0.5 ng/µL chymotrypsin (Roche, Basel, Switzerland; Promega, WI, USA) and incubated at 37°C for 5 h. The digested peptides were then dried in a vacuum freeze dryer. N-glycanase treatment was performed by dissolving the samples in 50 mM Tris-HCl (pH 8.0) and 10 nM ethylenediaminetetraacetic acid (EDTA) containing 3 units/μL PNGaseF (New England BioLabs, Inc. MA, USA) and incubated for 4 h at RT. The peptides were dried in a vacuum freeze dryer. For desalting, the samples were dissolved in 10% acetonitrile containing 0.1% formic acid and applied to a Ziptip (Merck Millipore) preconditioned with 95% acetonitrile containing 0.1% formic acid. After washing with 0.1% formic acid, bound peptides were eluted with 50% acetonitrile containing 0.1% formic acid.

### Liquid chromatography-tandem mass spectrometry analysis (LC-MS)

The peptides were analyzed using an Orbitrap Fusion Tribrid mass spectrometer (Thermo Fisher Scientific) coupled with an UltiMate3000 RSLC nano-LC system (Djonex Co.) and equipped with a nano-HPLC capillary column (150 mm x 75 μm; Nikkyo Technos, Tokyo, Japan). The LC gradient with 0.1% formic acid (solution A) and 0.1% formic acid in 90% acetonitrile (solution B) was set as follows: 5–100% B (0–45 min), 100–5% B (45–45.1 min), and 5% B (45.1–60 min). Data-dependent tandem mass spectrometry (MS) analysis was performed using a top-speed approach with a cycle time of 3 s.

The precursor ions were analyzed using an Orbitrap mass analyzer, whereas the fragment ions generated by the HCD were analyzed using a linear ion trap mass analyzer. Peak lists were generated using the extract msn function in Xcalibur 3.0.6 (Thermo Fisher Scientific) with the default parameters.

### Data analysis for mass spectrometry

Raw data were analyzed using Xcalibur 3.0.6 and Byonic Node (version 3.5.3). The parameter set included proteolytic cleavage sites at the C-termini of Lys/Arg/Phe/Trp/Tyr/Leu with the possibility of two missed cleavages, Cys carbamidomethylation as a fixed modification, Met oxidation as a common-1 modification, Asn/Gln deamidation as a common-1 modification, the O-glycan database (listed in Table S8) as a rare-1 modification, and 78 mammalian O-glycans as a rare-2 modification. The total common maximum was set to 3, and the total rare maximum was set to 2. MS/MS scoring was performed using the following parameters: precursor mass tolerance of 20 ppm, fragment mass tolerance of 20 ppm, and QTOF/HCD mode. Annotated peptides with a Byonic score > 200 and an abundance ratio > 3% in at least two triplicate samples were selected for analysis.

For semi-quantification, extracted ion chromatograms (EICs) were generated using the Xcalibur Qual browser by selecting the most abundant isotopic peaks with the following settings: smoothing enabled, Gaussian type at point 5, mass tolerance 5.0 ppm; and mass precision decimal 4, as described previously (43).

### Immunoblotting

Samples were separated by sodium dodecyl sulfate-polyacrylamide gel electrophoresis and transferred onto polyvinylidene difluoride membranes (Merck Millipore). The membranes were incubated with 3% skim milk dissolved in phosphate-buffered containing 0.05% Tween 20 (PBS-T). For detection, membranes were incubated with the indicated primary antibodies diluted in 3% skim milk in PBS-T at RT for 1h. This was followed by incubation with the appropriate horseradish peroxidase (HRP)-conjugated secondary antibodies at RT for 1h. The bands were visualized using Immobilon Western Chemiluminescent HRP Substrate (Merck Millipore) exposed to X-ray film (Fujifilm, Tokyo, Japan) or detected using the iBright FL1500 Imaging System (Invitrogen).

### Analysis of FLAG-DLK1 turnover using cycloheximide

Cells grown in a 10 cm dish were transfected with plasmids encoding FLAG-DLK1 using PEI-Max. The following day, the cells were harvested using trypsin/EDTA and divided into six 3.5 cm dishes. The next day, the cells were incubated in complete culture medium containing 20 μg/mL cycloheximide at 37°C for the indicated times.

### Secretion Assay for DLK1-ECD

Cells grown in 6-well plates were transiently transfected with plasmids encoding DLK1-ECD using PEI-Max. On the following day, the medium was replaced with Opti-MEM. After three days of culture, the medium was collected and sequentially centrifuged at 4°C at increasing speeds, including 900 *× g* for 3 min (to remove dead cells and cell debris) and 17,000 *× g* for 15 min (to remove insoluble substances). Cell lysates were prepared as described previously.

### Transport Assay for Reporter DLK1

Cells grown in 6-well plates were transiently transfected with plasmids expressing hook proteins and the Reporter DLK1. The next day, the medium was replaced with the complete culture medium containing 1 μg/μL doxycycline. After incubation for one day, the cells were washed twice with phosphate-buffered saline (PBS) and harvested using trypsin/ethylenediaminetetraacetic acid (EDTA). After washing with HBSS+, cells were incubated in 50 μL of DMEM containing 200 μM D-biotin (Merck), 20 mM Tris-HCl (pH 7.4), and 10% FBS for the indicated times. After washing with 1% BSA-HBSS+, the cells were stained with an anti-DLK1 antibody, followed by PE-conjugated anti-mouse IgG staining, and subsequent flow cytometry analysis. The GFP-positive cells were gated at a constant intensity for all samples. The mean fluorescence intensity (MFI) of DLK1 staining in control HEK293 cells after 60 min of chase was set as 100% transport.

### Immunoprecipitation

Cells grown in 6-well plates were transiently transfected with plasmids expressing hook proteins and the Reporter DLK1. The next day, the medium was replaced with the complete medium containing 1 μg/μL doxycycline. After incubation for one day, cell lysates were prepared as described above and subjected to immunoprecipitation with anti-DYKDDDDK-tag antibody clone 1E6 or mouse IgG anti-HA antibody clone 4B2 using Protein G Dynabeads (Thermo Fisher Scientific). The beads (bound fraction) were washed three times with lysis buffer. To detect the Reporter DLK1 and Hook proteins, the anti-FLAG antibody clone 15 and anti-HA antibody clone C29F4 were used.

### PNGase F treatment

FLAG-DLK1 or Reporter DLK1 was collected from cell lysates using a DDDDK-tagged protein purification gel (clone FLA, MBL) at 4°C overnight. The gels were incubated with or without 25 units/μL of PNGase F (NEB) at 37°C for 1 h under denatured conditions according to the instructions by the manufacturer.

### Immunofluorescence microscopy

To analyze the ER localization of the Reporter DLK1^del^, BFP-KDEL was co-expressed with the Reporter DLK1^del^ in HEK293 cells. After 6 h, the transfected cells were replated on a Cellmatrix Type I-A-coated glass slip (Nitta Gelatin. Inc, Osaka, Japan). After overnight culture, 1 μg/mL doxycycline was added to induce Reporter DLK1^del^ expression. At 24 h after induction, cells were fixed with 4% paraformaldehyde (PFA) for 10 min. After three washes with PBS, the cells were permeabilized with 3% BSA and 0.05% Triton X-100 in PBS for 1 h and then mounted in TOPRO3 (Thermo Fisher Scientific)-containing mounting medium.

To observe Reporter DLK1^del^ trafficking, cells were incubated with a complete culture medium containing 1 μg/mL doxycycline for 24 h. Then, D-biotin was added and chased at the indicated time points. The cells were fixed and permeabilized as described above. The cells were stained with anti-GM130 antibodies diluted in 0.05% Triton X-100 and 3% BSA in PBS for 1 h at RT. After washing three times with PBS, the cells were mounted in a TOPRO3-containing mounting medium. Confocal microscopy images were obtained using a Nikon A1-Rsi confocal microscope equipped with a Plan-Apo 60X/1.40 oil immersion objective lens.

## Abbreviations

DLK1: Delta-like 1 homolog
EGF: Epidermal growth factor-like
ER: Endoplasmic reticulum
O-Fuc: O-Fucose
O-Glc: O-Glucose
O-GlcNAc: O-N-Acetyl-glucosamine
ECD: Ectodomain
PEI: Polyethyleneimine
RT: Room temperature
PTM: Post-translational modification
O-GalNAc: O-N-Acetyl-galactosamine
RUSH: Retention using selective hooks

## Data Availability

All raw and processed LC-MS/MS data files were deposited in jPOST (Project ID: JPST003303).

## Author contributions

YT: Conceptualization, Visualization, Investigation, Validation, Formal analysis, data curation, resources, writing, editing, funding acquisition, writing–original draft, writing–review, and editing; YoT: Data curation, formal analysis; YK: Investigation, Writing–review, and editing; NT: Investigation, Data curation; EU: Investigation; WF: Investigation; SG: Investigation, Resources; HT: Supervision, Writing–review, and editing; TO: Conceptualization, Visualization, Funding acquisition, project administration, Supervision, Writing–original draft, writing–review, and editing.

## Acknowledgments

We thank Hiroyuki Kaji and Kentaro Taki (Nagoya University) for assistance with LC-MS/MS analysis; Yusuke Maeda and Taroh Kinoshita (Osaka University) for the pCX-bsr-human_ManII-G4S2×8-EGFP-SBP and pCMV-hyPBase-teton-P2A-BSD plasmids; Chika Saito, Rashu Barua, Mitsutaka Ogawa, and Kenta Sakuma (Nagoya University) for assistance with the initial phase of the project. This work was conducted under the Collaborative Open Research Program to promote the Human Glycome Atlas Project (HGA) as strategic interdisciplinary research in the J-GlycoNet cooperative network that is accredited by the Minister of Education, Culture, Sports, Science, and Technology, MEXT, Japan, as a Joint Usage/Research Center.

