## Supporting information for "Retention of DLK1 in the endoplasmic reticulum identifies roles for EGF domain-specific O-glycans in the secretory pathway"

for

**Running title:**
Role of atypical O-glycan of DLK1 in the secretory pathway

**Keywords:**
O-glycan, DLK1, EGF, glycoproteomics, transport

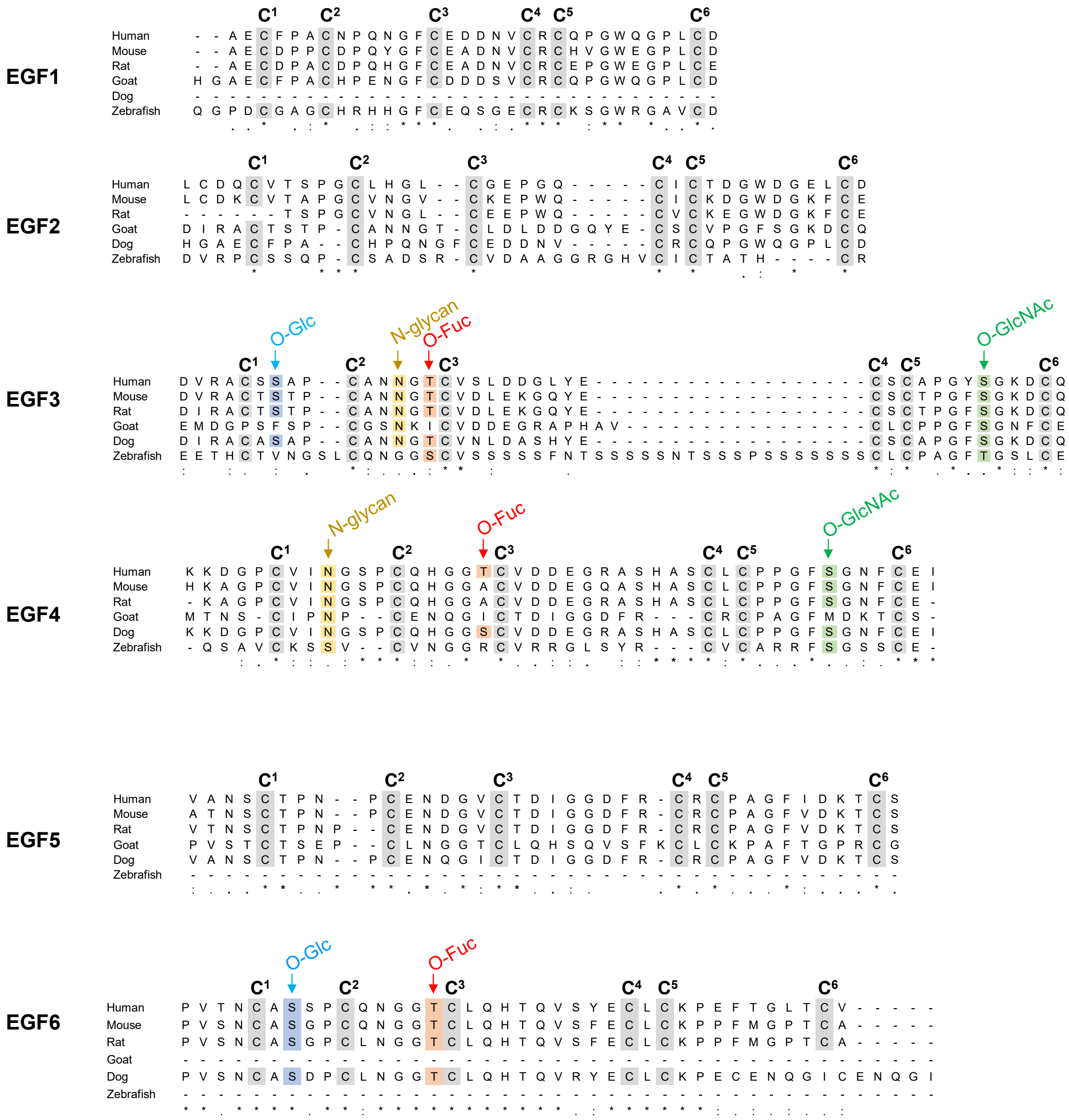

**Supporting Figure S1. Alignment of amino acid sequences of DLK1 from different species.**
Multiple sequence alignments were performed using ClustalW 2.1. Human DLK1, *Homo sapiens* (NP_003827.4); mouse DLK1, *Mus musculus* (GenBank accession number AAH52159. 1); rat DLK1, *Rattus norvegicus* (NP_446196.1); goat DLK1, *Capra hircus* (NP_001301141.1); dog DLK1, *Canis lupus familiaris* (XP_038528912.1); zebrafish DLK1, *Danio rerio* (XP_005173621.1). The arrow indicates the consensus amino acid sequence for each glycosylation site. Gray highlights indicate six conserved cysteine residues comprising an EGF repeat.

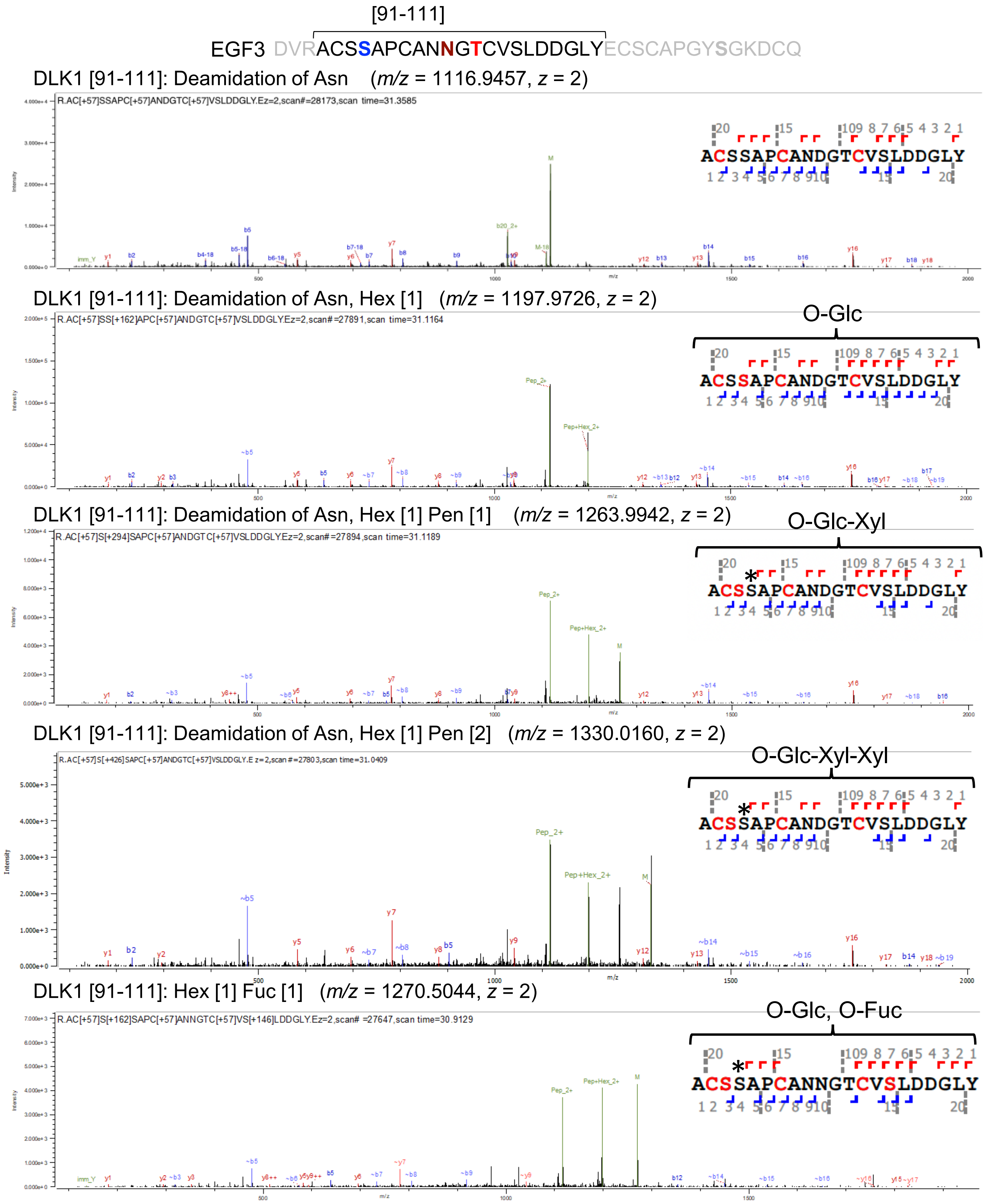


**Supporting Figure S2. In tandem mass spectrometry (MS/MS) spectra of the DLK1[91-111] fragment derived from DLK1-ECD.**

Byonic-assisted annotation of higher-energy collisional dissociation in tandem mass spectrometry (HCD-MS/MS) spectra showing proteolytic fragments of DLK1[91-111] modified with N-glycans and/or indicated O-glycans. DLK1-ECD was purified from the culture medium of transfected HEK293T cells and was digested with trypsin and chymotrypsin. Proteolytic fragments were treated with PNGase to remove N-glycans. The resulting conversion of asparagine (N) to aspartic acid (D) (deamidation) and O-glycan modifications were analyzed using liquid chromatography in tandem mass spectrometry (LC-MS/MS). The amino acid residues located at the putative O-glucose (O-Glc), O-fucose (O-Fuc), O-GlcNAc, and N-glycan modification sites are indicated in blue, red, green, and brown, respectively, at the top of the figure. The modification site for the proposed O-glycan structure in the inset is not fully supported by the HCD-MS/MS data, and in some cases, does not agree with the predicted modification site (indicated by an asterisk).

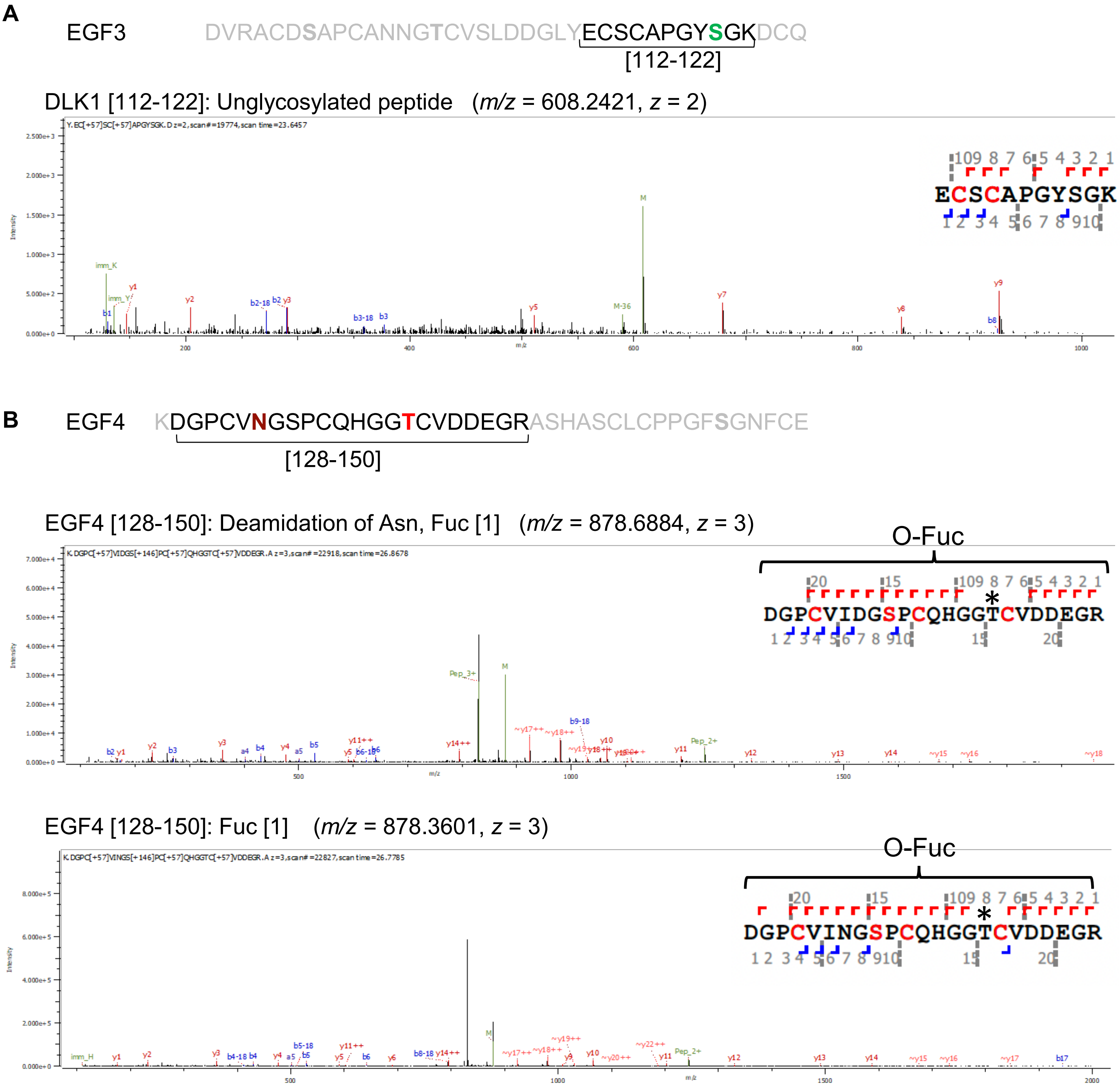


**Supporting Figure S3. MS/MS spectra of DLK1[112-122] and DLK1[128-150] fragments from DLK1-ECD.** Byonic-assisted annotation of HCD-MS/MS spectra showing proteolytic fragments of DLK1[112-122] (A) or DLK1[128-150] (B) modified with N-glycans and/or the indicated O-glycans. DLK1-ECD was purified from the culture medium of the transfected HEK293T cells and digested with trypsin and chymotrypsin. Proteolytic fragments were treated with PNGase to remove N-glycans. The resulting conversion of asparagine (N) to aspartic acid (D) (deamidation) and O-glycan modifications were analyzed using LC-MS/MS. Amino acid residues located at the putative modification sites for O-Fuc, O-GlcNAc, and N-glycan are indicated in red, green and brown, respectively, at the top of the panel. The modification site for the proposed O-glycan structure in the inset is not fully supported by the HCD-MS/MS data, and in some cases, it does not agree with the predicted modification site (indicated by an asterisk).

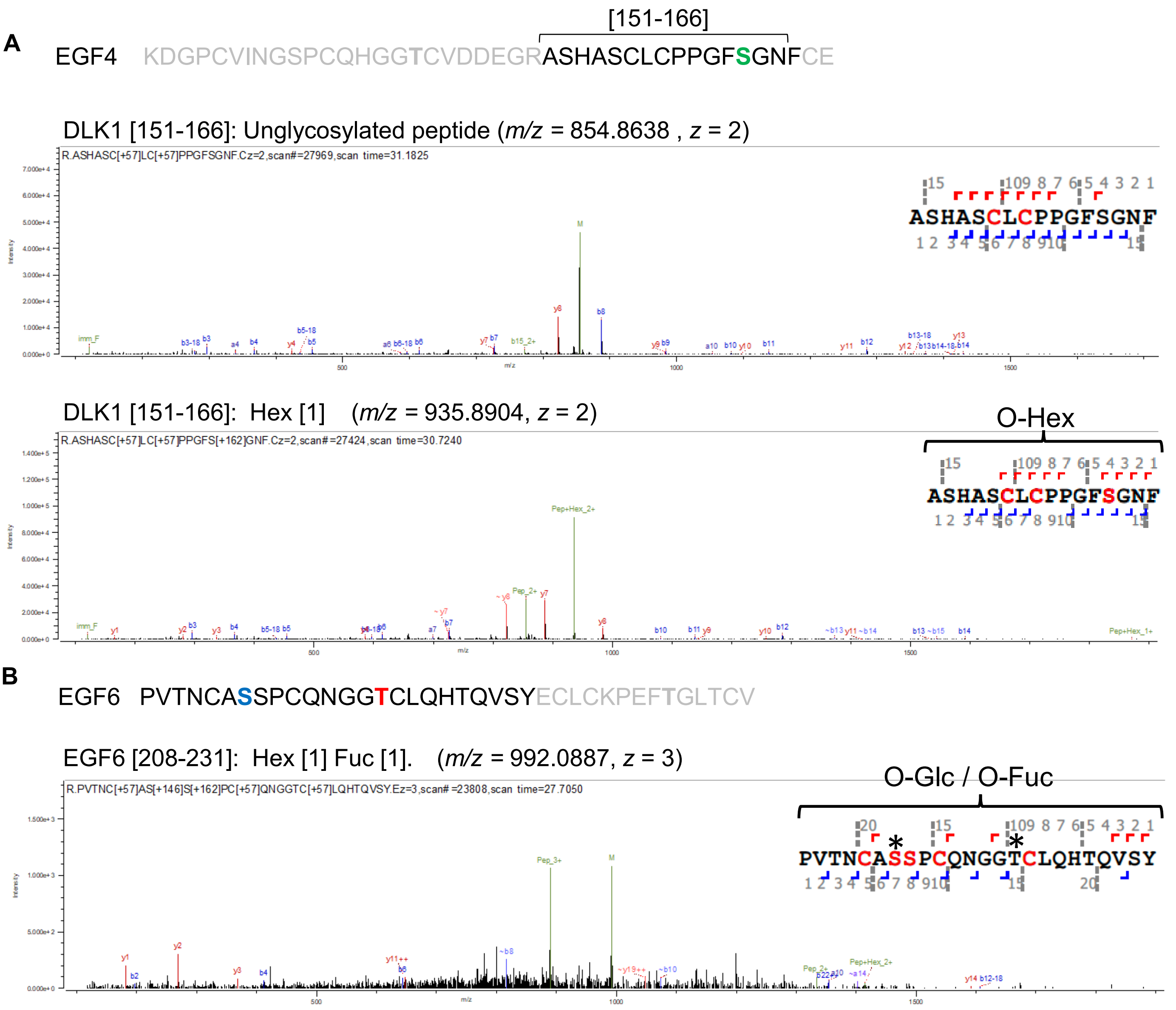


**Supporting Figure S4. MS/MS spectra of DLK1[151-166] and DLK1[208-231] fragments from DLK1-ECD.** Byonic-assisted annotation of HCD-MS/MS spectra showing proteolytic fragments of DLK1[151-166] (A) or DLK1[208-231] (B) modified with or without the indicated O-glycans. DLK1-ECD was purified from the culture medium of transfected HEK293T cells and was digested with trypsin and chymotrypsin. Proteolytic fragments were treated with PNGase to remove N-glycans and O-glycan modifications were analyzed using LC-MS/MS. Amino acid residues located at the putative modification sites for O-Glc, O-Fuc, and O-GlcNAc are indicated in blue, red, and green, respectively, at the top of the panel. The modification site for the proposed O-glycan structure in the inset is not fully supported by the HCD-MS/MS data, and in some cases, does not agree with the predicted modification site (indicated by an asterisk).

### DLK1-ECD expressed in wild type HEK293T cells

DLK1 [267-283]
RLTPGVHELPVQQPEHR + HexNAc [1] Hex [1] NeuAc [1]
*m/z* = 663.0806, *z* = 4

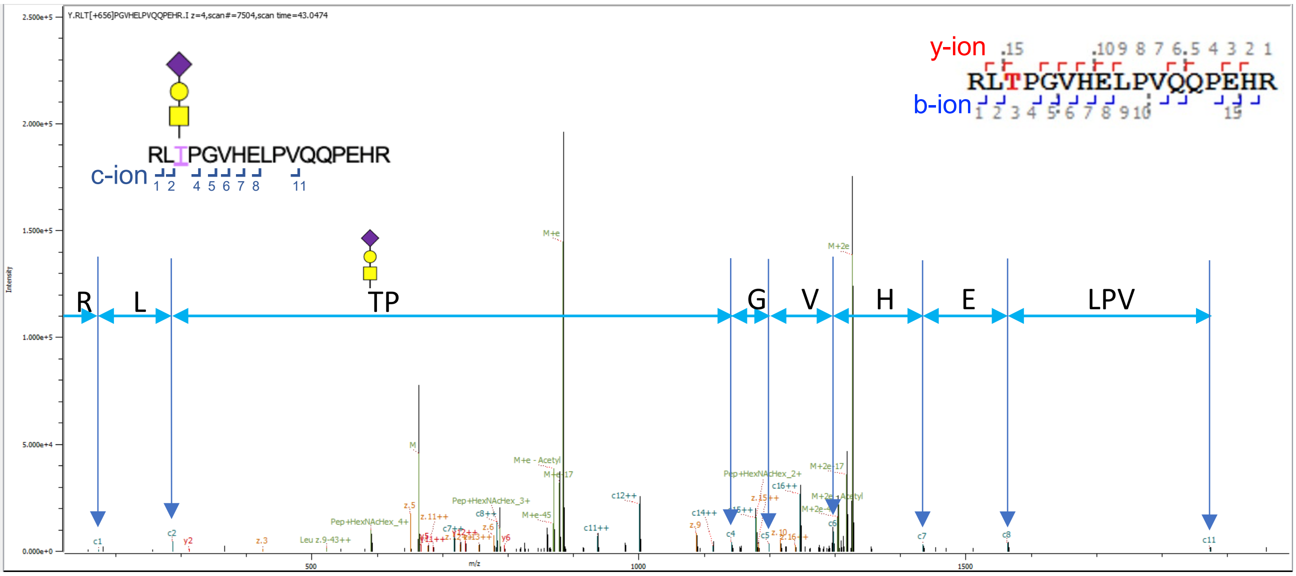


**Supporting Figure S5. MS/MS spectra of DLK1[268-283] fragment derived from DLK1-ECD.**
Byonic-assisted annotation of HCD-aided electron transfer dissociation (EThcD) D-MS/MS spectra showing proteolytic fragments of DLK1[268-283] modified with O-GalNAc glycans. DLK1-ECD was purified from the culture medium of transfected HEK293T cells and digested with trypsin and chymotrypsin. Proteolytic fragments were treated with PNGase to remove N-glycans, and O-glycan modifications were analyzed using LC-MS/MS. Amino acid residues modified by O-GalNAc glycans are shown in the insets.

**Experimental Procedures For Supporting Figure S5**
EThcD fragmentation was performed by LC-MS/MS using an Orbitrap Fusion Eclipse Tribrid mass spectrometer (Thermo Fisher Scientific, Waltham, MA, USA) equipped with an nLC system Ultimate 3000 (Thermo Fisher Scientific) and an octadecylsilyl tip column (inside diameter 75 µm, 150 mm, bead size: 3 µm, Nikkyo Technos, Tokyo, Japan). Mobile phases A and B contained 0.1% formic acid in double distilled water and 0.1% formic acid in acetonitrile, respectively (Kanto Chemical, Tokyo, Japan). Peptides were dissolved in 0.1% formic acid in Milli-Q water. Peptides were separated using a 3–35% acetonitrile gradient for 90 min. The mass spectrometer was operated in the positive and data-dependent acquisition modes. Master scan MS spectra (m/z 377–2000) were acquired using an Orbitrap mass analyzer with 120 K resolution and a Normalized AGC target with the following parameters: 100%, AGC target: 4E5, and RF lens: 30%. To obtain the MS/MS spectra, the precursors were collected by a filter charge state: z=2–7; dynamic exclusion: 15 s; intensity threshold: max= 1E+20, min=5E4; and precursor selection range: m/z 400–2,000 with a cycle time of 3 s. The selected precursors were fragmented using HCD and EThcD with a maximum cycle time of 3 s. HCD was performed according to a stepped collision method (20, 30, and 40%) and detected using an Orbitrap at 15 K resolution with a scan range of m/z 135–2000 and a 100% AGC target. EThcD was performed using a charge-dependent ETD at a maximum injection time of 300 ms and an AGC target of 5E4. MS/MS spectra were obtained using an Orbitrap analyzer at 15 K resolution with a scan range of m/z 135–2,000.

**A** DLK1-ECD expressed in *EOGT*-deficient cells

EGF4: KDGPCVINGSPCQHGGTCVDDEGRASHASCLCPPGF**S**GNFCE

[151-166]

DLK1[151-166]: Unglycosylated peptide ($m/z$ = 854.8641, $z$ = 2)

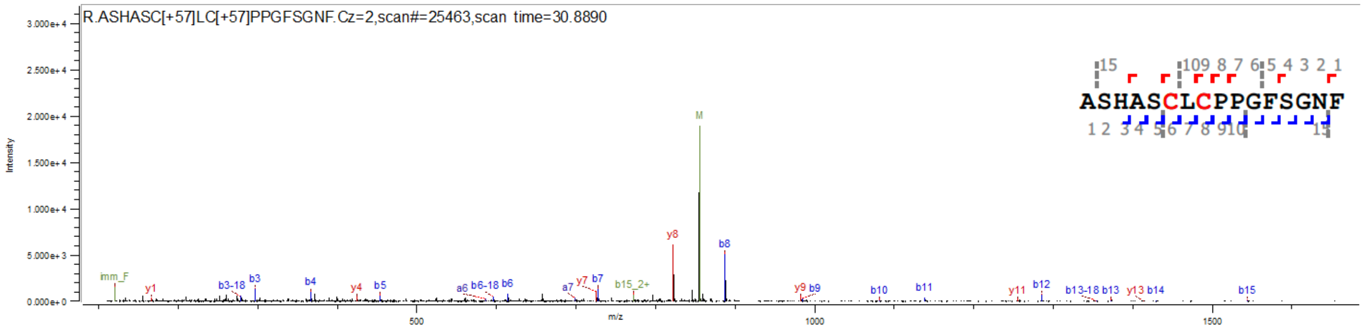


**B** DLK1-ECD[S163A] expressed in HEK293T cells

EGF4: KDGPCVINGSPCQHGGTCVDDEGRASHASCLCPPGF**A**GNFCE

[151-166]

DLK1[S163A] [151-166]: Unglycosylated peptide ($m/z$ = 846.8666, $z$ = 2)

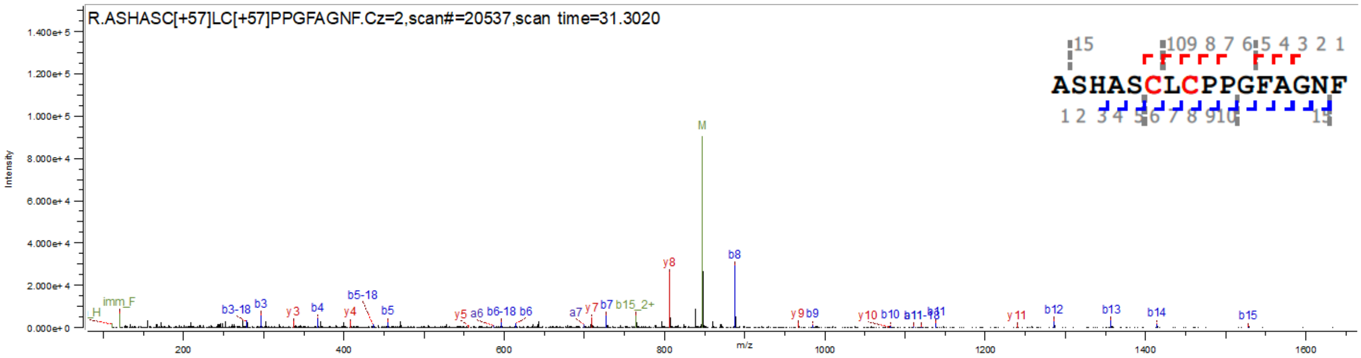


**Supporting Figure S6. MS/MS spectra of DLK1-ECD produced in *EOGT*-deficient HEK293T cells and the S163A mutant.**

Byonic-assisted annotation of HCD-MS/MS spectra showing proteolytic fragments of DLK1[151-166] expressed in *EOGT*-deficient HEK293T cells (A) and S163A mutant produced in *EOGT*-deficient HEK293T cells (B). Purified proteins were digested with trypsin and chymotrypsin. Proteolytic fragments were treated with PNGase to remove N-glycans, and O-glycan modifications were analyzed using LC-MS/MS. The amino acid residue located at the putative O-GlcNAc modification site is indicated in green at the top of the panel (A).

**DLK1-ECD expressed in wild type HEK293T cells**

**DLK1[249-257]**

DLK1 [249-257]: HexNAc [3] Hex [1] NeuAc [1] (*m/z* = 687.9856, *z* = 3)

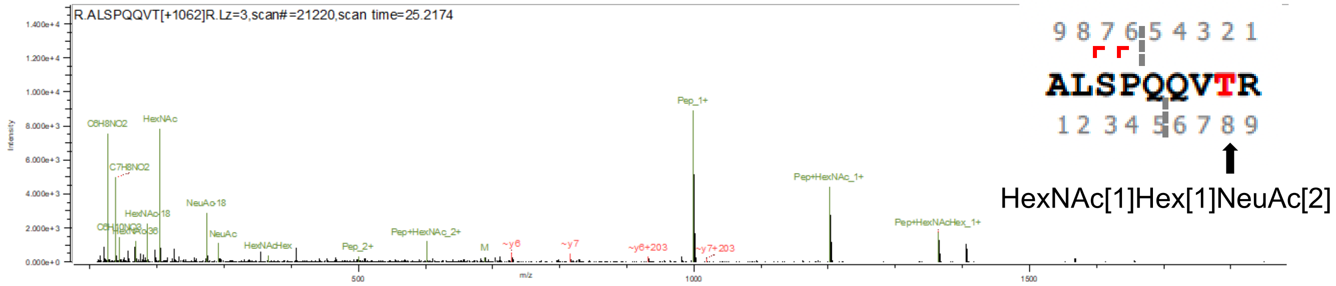


DLK1 [249-257]: HexNAc [1] Hex [1] NeuAc [2] (*m/z* = 649.6325, *z* = 3)

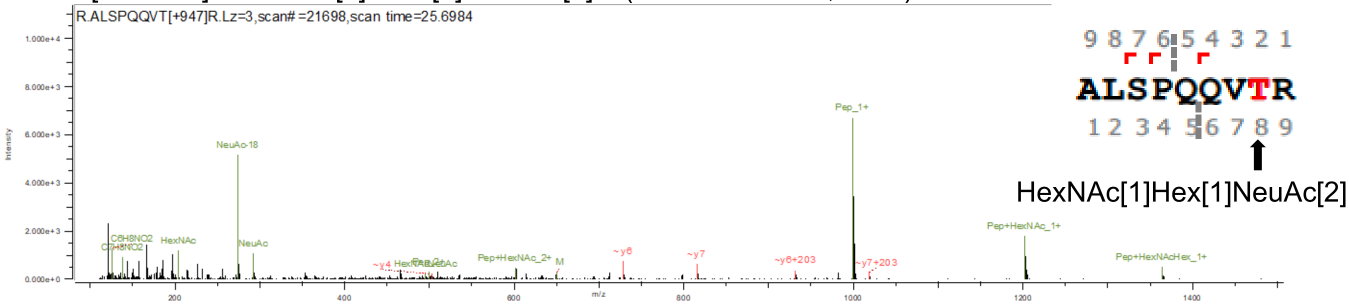


DLK1 [249-257]: HexNAc [1] Hex [1] NeuAc [2] (*m/z* = 973.9446, *z* = 2)

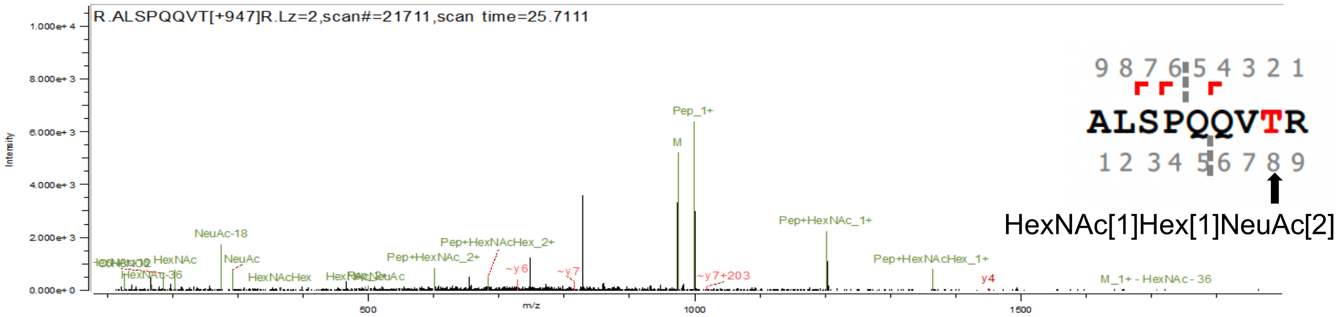


DLK1 [249-257]: HexNAc [2] (*m/z* = 703.3618, *z* = 2)

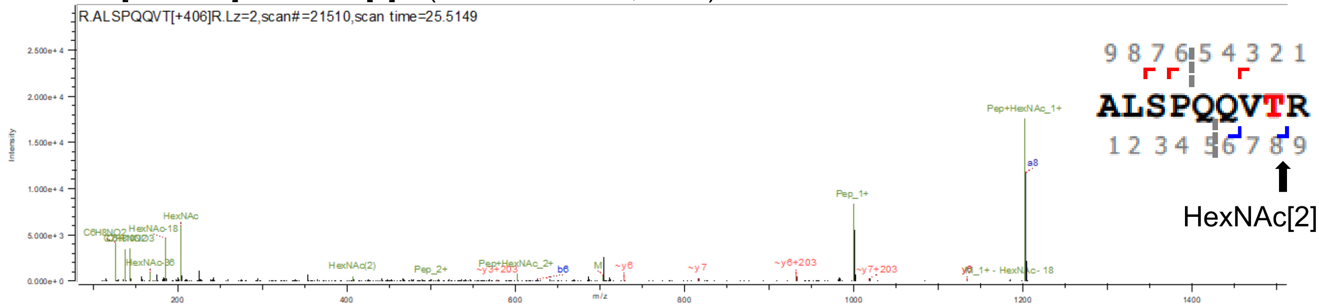


DLK1 [249-257]: Unglycosylated peptide (*m/z* = 500.2823, *z* = 2)

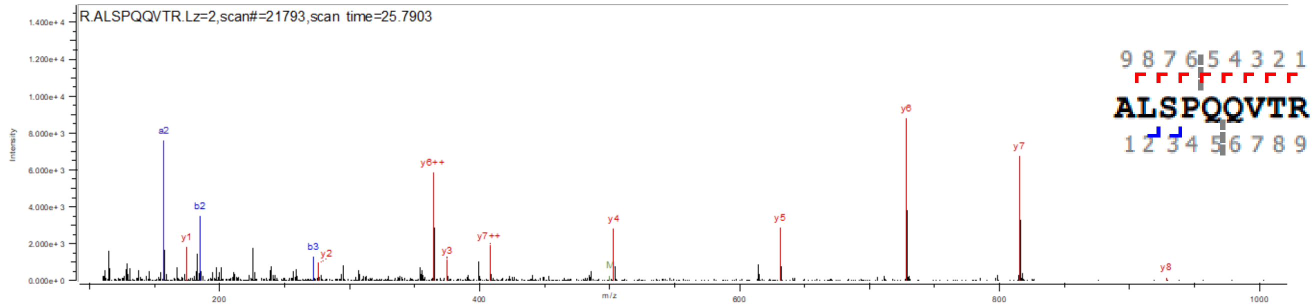


**Supporting Figure S7. MS/MS spectra of the DLK1[249-257] fragment derived from DLK1-ECD.** Byonic-assisted annotation of HCD-MS/MS spectra showing proteolytic fragments of DLK1[249-257] modified with O-GalNAc glycans. DLK1-ECD was purified from the culture medium of transfected HEK293T cells and digested with trypsin and chymotrypsin. Proteolytic fragments were treated with PNGase to remove N-glycans, and O-glycan modifications were analyzed using LC-MS/MS. Amino acid residues modified by O-GalNAc glycans are indicated by arrows in the insets.

### DLK1-ECD expressed in wild type HEK293T cells

DLK1 [268-283]: HexNAc [3] Hex [1] NeuAc [1] (*m/z* = 725.5950 *z* = 4)

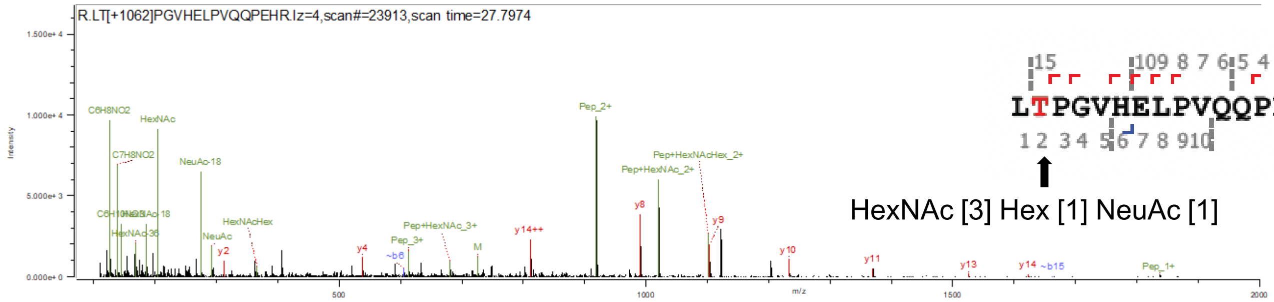


DLK1 [268-283]: HexNAc [1] Hex [1] NeuAc [1] (*m/z* = 624.0555, *z* = 4)

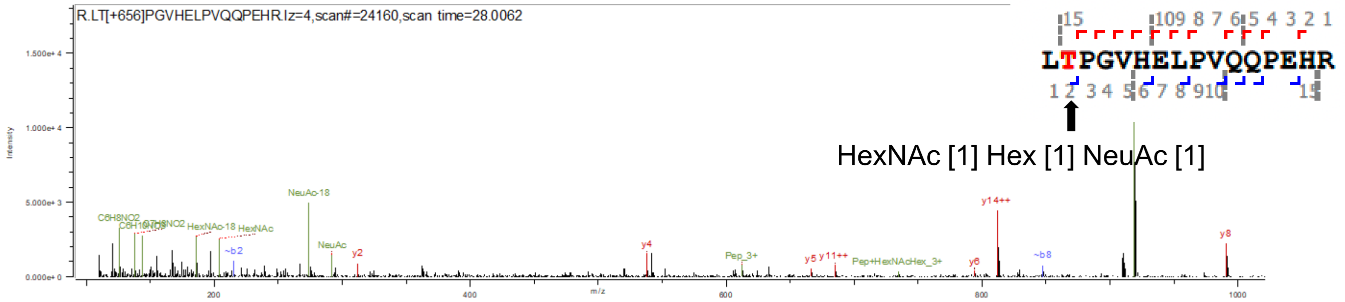


DLK1 [268-283]: HexNAc [2] (*m/z* = 561.5368, *z* = 4)

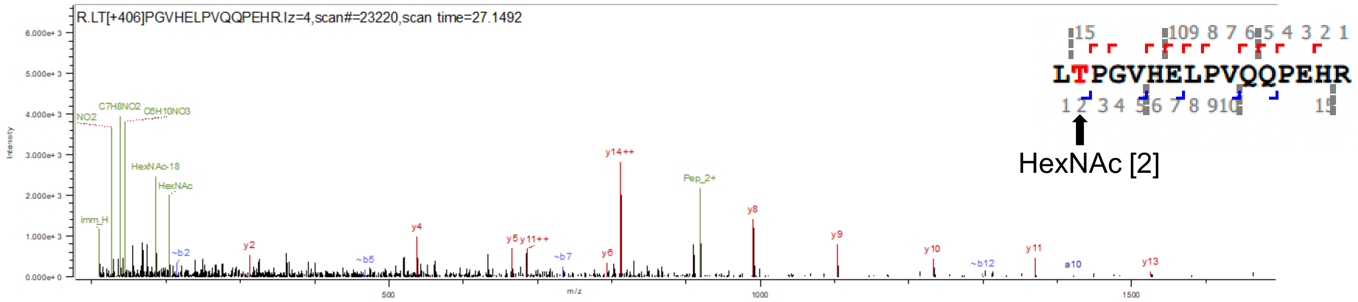


DLK1 [268-283]: Un-glycosylated (*m/z* = 612.9954, *z* = 3)

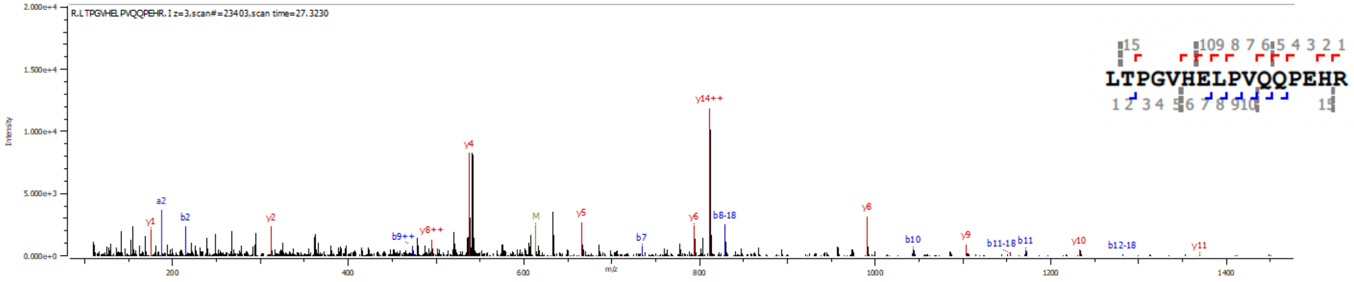


DLK1 [268-283]: Un-glycosylated (*m/z* = 459.9986, *z* = 4)

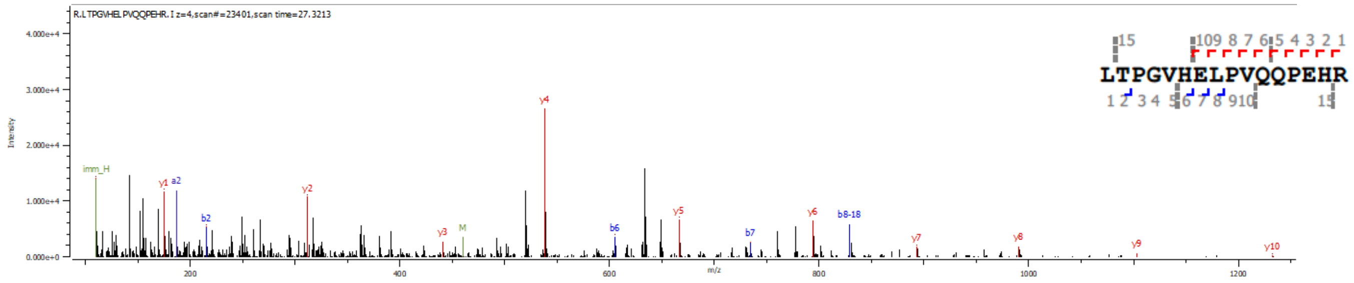


**Supporting Figure S8. MS/MS spectra of the DLK1[268-283] fragment derived from DLK1-ECD.** Byonic-assisted annotation of HCD-MS/MS spectra showing proteolytic fragments of DLK1[268-283] modified with O-GalNAc glycans. DLK1-ECD was purified from the culture medium of transfected HEK293T cells and digested with trypsin and chymotrypsin. Proteolytic fragments were treated with PNGase to remove N-glycans, and O-glycan modifications were analyzed using LC-MS/MS. The amino acid residues modified by O-GalNAc glycans are indicated by arrows in the insets.

**Reporter DLK1 expressed in wild type HEK293 cells**

DLK1 [249-257]: HexNAc [1]     (m/z = 601.8228, *z* = 2)

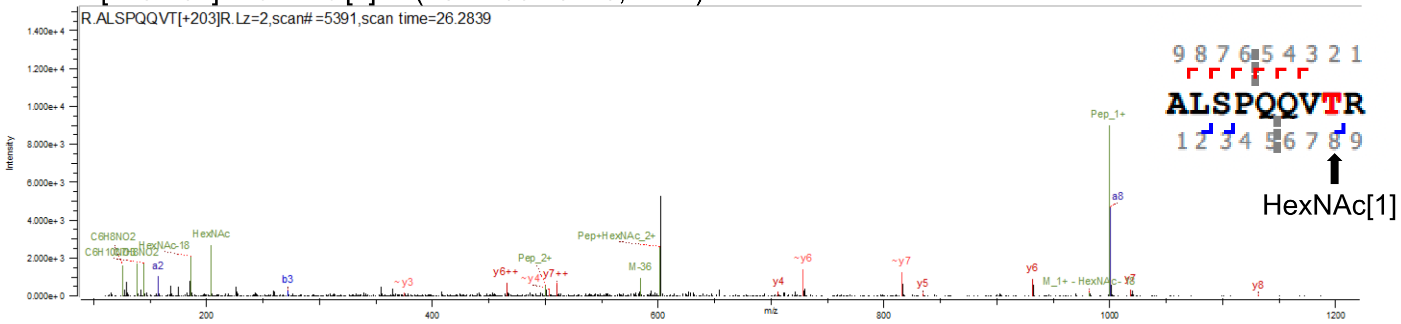


DLK1 [249-257]: Unglycosylated peptide    (*m/z* = 500.2829, *z* = 2)

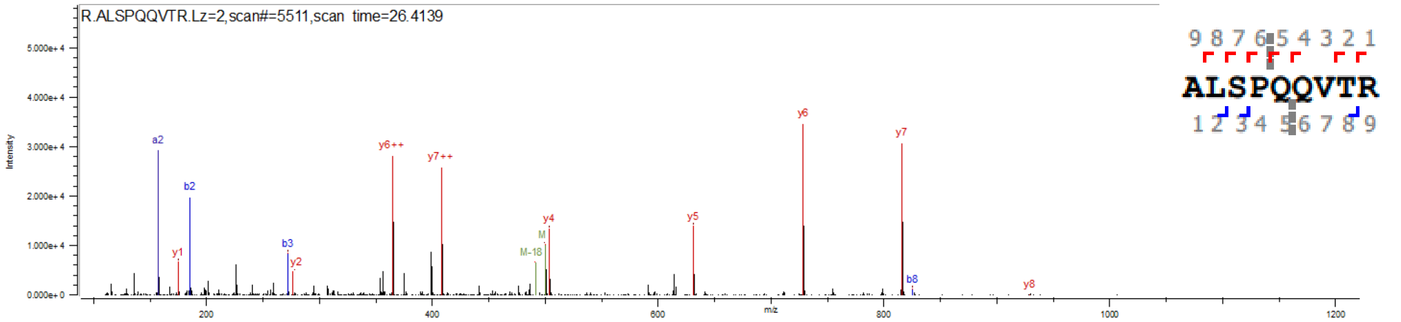


**Supporting Figure S9. MS/MS spectra of the DLK1[249-257] fragment derived from reporter DLK1.** Byonic-assisted annotation of HCD-MS/MS spectra showing proteolytic fragments of DLK1[249-257] modified with O-GalNAc glycans. The reporter DLK1 was prepared from HEK293T cells and digested with trypsin and chymotrypsin. Proteolytic fragments were treated with PNGase to remove N-glycans, and O-glycan modification was analyzed using LC-MS/MS. The amino acid residues modified by O-GalNAc glycans are indicated by arrows in the insets.

### Reporter DLK1 expressed in wild type HEK293 cells

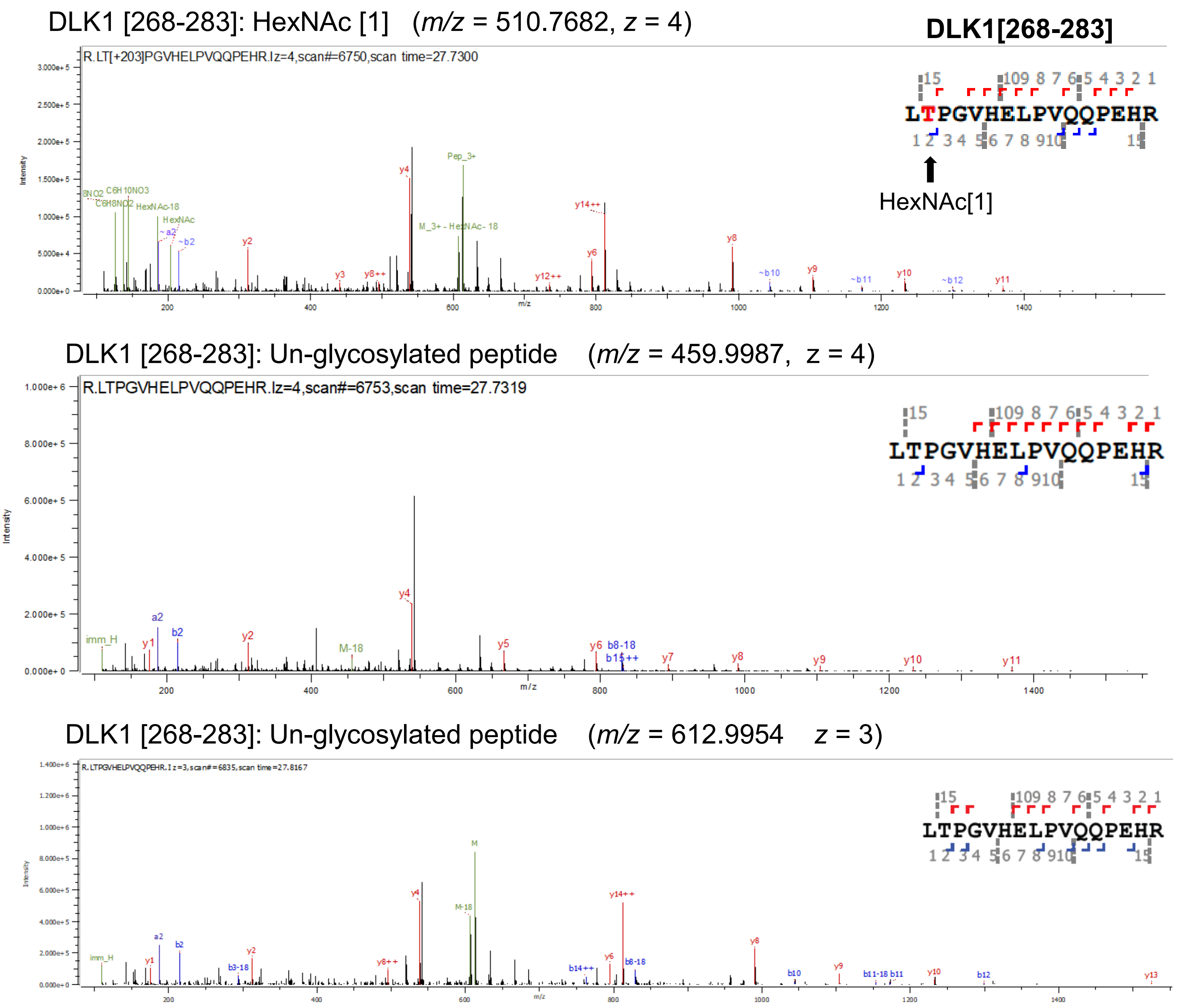


**Supporting Figure S10. MS/MS spectra of the DLK1[268-283] fragment derived from reporter DLK1.** Byonic-assisted annotation of HCD-MS/MS spectra showing proteolytic fragments of DLK1[249-257] modified with O-GalNAc glycans. The reporter DLK1 was prepared from HEK293T cells and digested with trypsin and chymotrypsin. Proteolytic fragments were treated with PNGase to remove N-glycans, and O-glycan modification was analyzed using LC-MS/MS. The amino acid residues modified by O-GalNAc glycans are indicated by arrows in the insets.

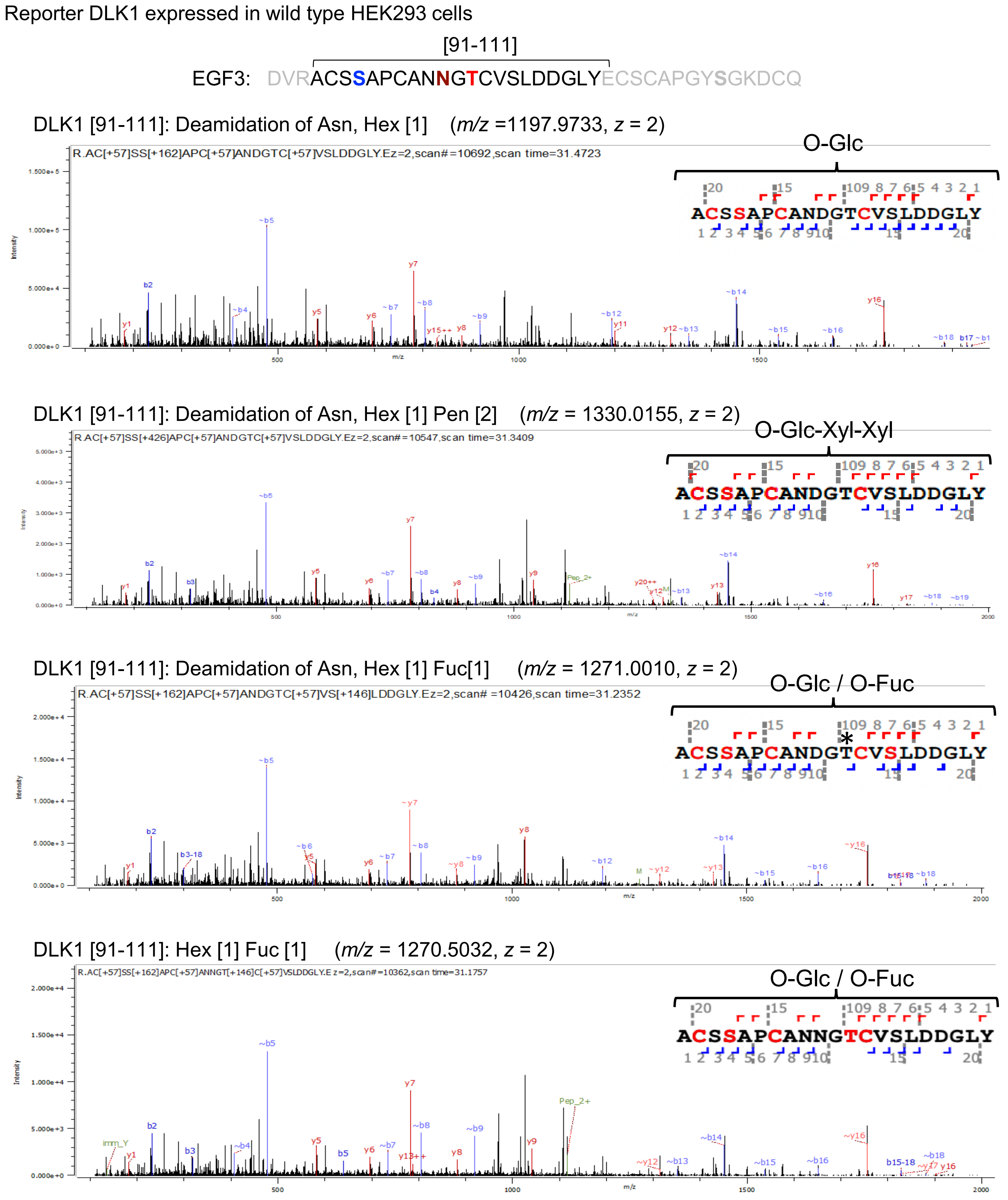


**Supporting Figure S11. MS/MS spectra of the DLK1[91-111] fragment of reporter DLK1 expressed in wild-type HEK293 cells.**

Byonic-assisted annotation of HCD-MS/MS spectra showing proteolytic fragments of DLK1[91-111] modified with N-glycans and/or the indicated O-glycans. The reporter DLK1 was prepared from HEK293 cells and digested with trypsin and chymotrypsin. Proteolytic fragments were treated with PNGase to remove N-glycans. The resulting conversion of asparagine (N) to aspartic acid (D) (deamidation) and O-glycan modifications were analyzed using LC-MS/MS. The amino acid residues located at the putative O-Glc, O-Fuc, and N-glycan modification sites are indicated in blue, red, and brown, respectively, at the top of the figure. The modification site for the proposed O-glycan structure in the inset is not fully supported by the HCD-MS/MS data, and in some cases, it does not agree with the predicted modification site (indicated by an asterisk).

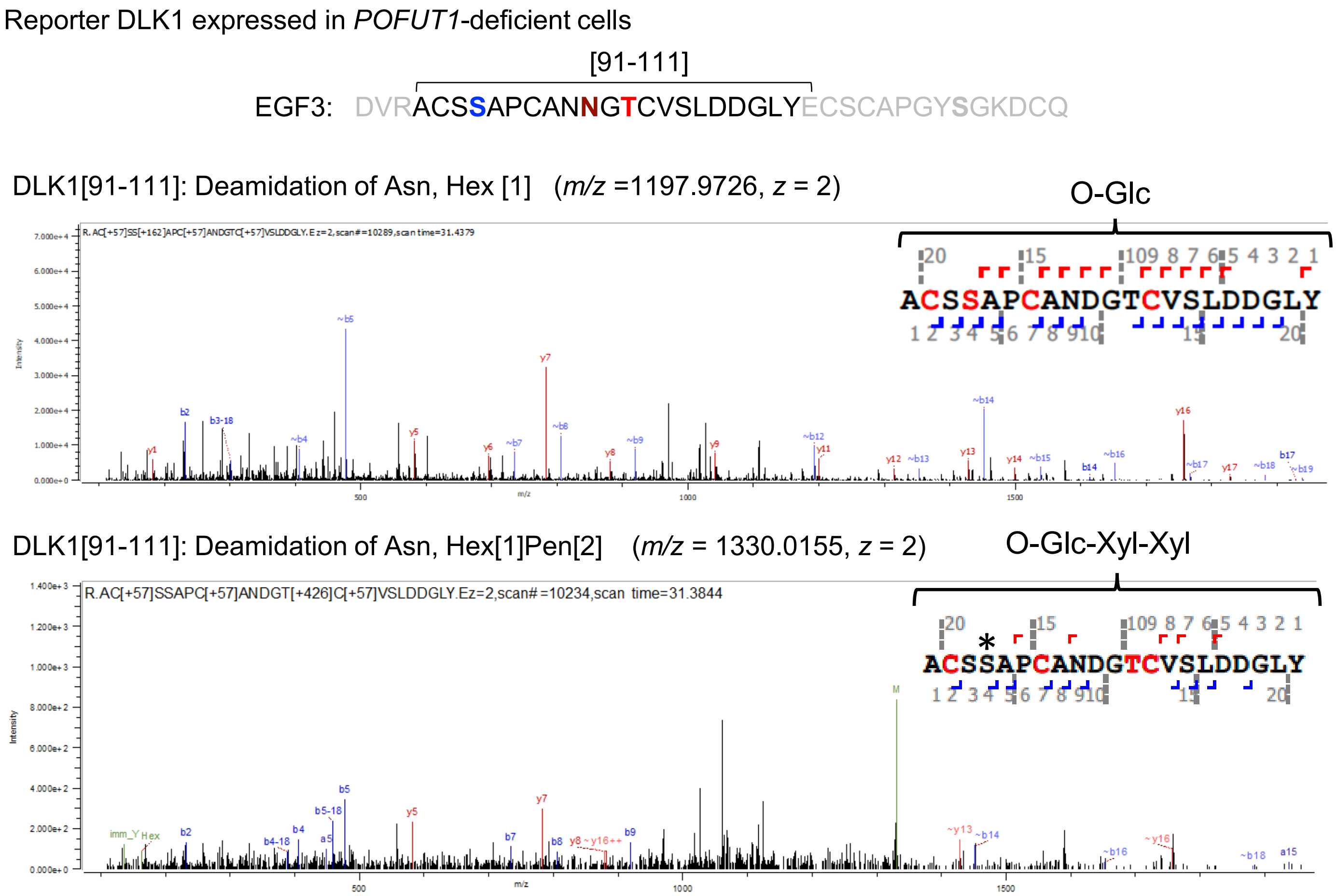


**Supporting Figure S12. MS/MS spectra of the DLK1[91-111] fragment of reporter DLK1 expressed in *POFUT1*-deficient cells.**

Byonic-assisted annotation of HCD-MS/MS spectra showed proteolytic fragments of DLK1[91-111] modified with N-glycans and the indicated O-glycans. The reporter DLK1 was prepared from *POFUT1*-deficient HEK293 cells and digested with trypsin and chymotrypsin. Proteolytic fragments were treated with PNGase to remove N-glycans. The resulting conversion of asparagine (N) to aspartic acid (D) (deamidation) and O-glycan modifications were analyzed using LC-MS/MS. The amino acid residues located at the putative O-Glc, O-Fuc, and N-glycan modification sites are indicated in blue, red, and brown, respectively, at the top of the figure. The modification site for the proposed O-glycan structure in the inset is not fully supported by the HCD-MS/MS data, and in some cases, it does not agree with the predicted modification site (indicated by an asterisk).

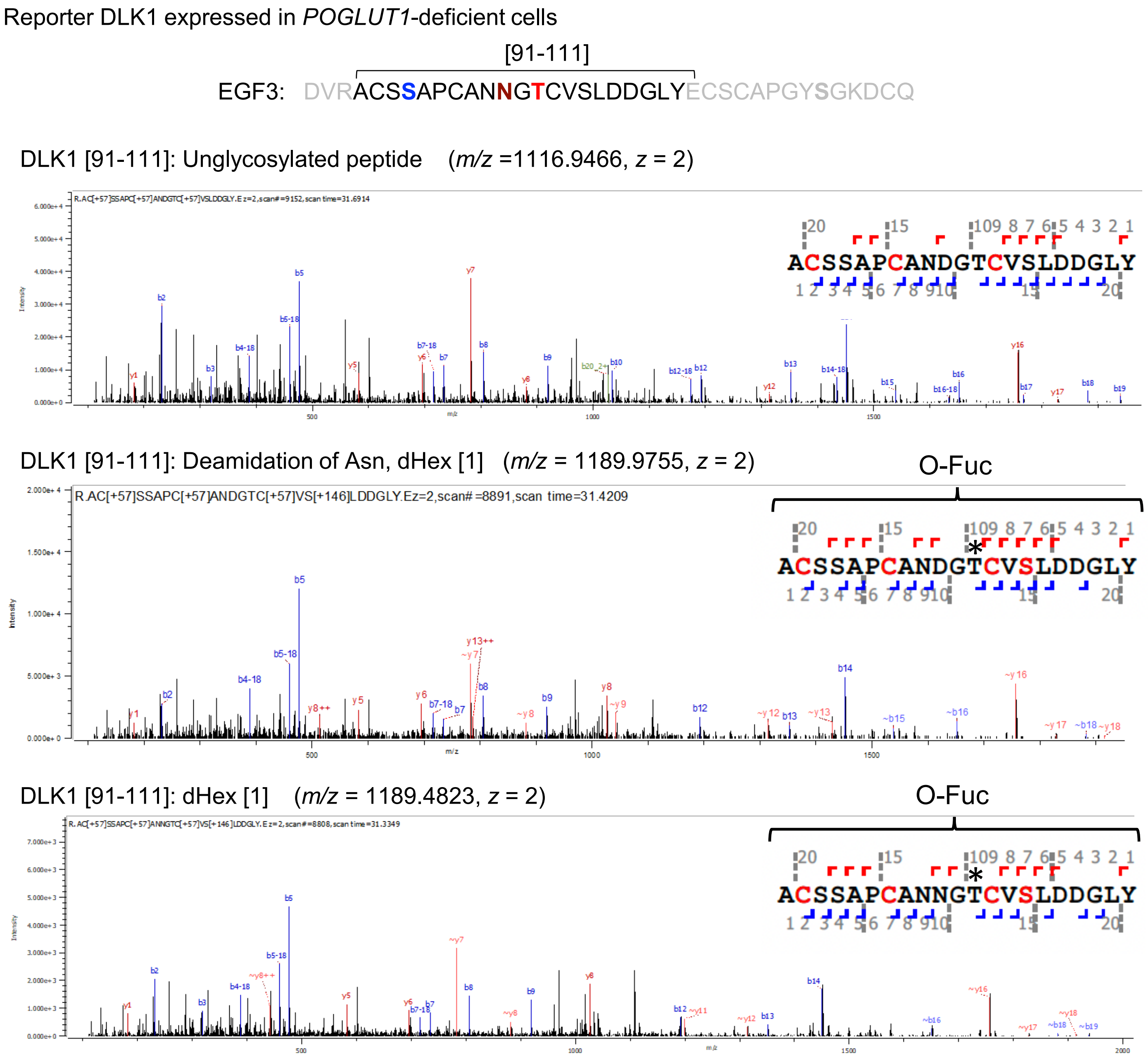


**Supporting Figure S13. MS/MS spectra of the DLK1[91-111] fragment of reporter DLK1 expressed in *POGLUT1*-deficient cells.**

Byonic-assisted annotation of HCD-MS/MS spectra showing proteolytic fragments of DLK1[91-111] modified with N-glycans and/or indicated O-glycans. Reporter DLK1 was purified from the culture medium of transfected *POGLUT1*-deficient HEK293 cells and digested with trypsin and chymotrypsin. Proteolytic fragments were treated with PNGase to remove N-glycans. The resulting conversion of asparagine (N) to aspartic acid (D) (deamidation) and O-glycan modifications were analyzed using LC-MS/MS. The amino acid residues located at the putative O-Glc, O-Fuc, and N-glycan modification sites are indicated in blue, red, and brown, respectively, at the top of the figure. The modification site for the proposed O-glycan structure in the inset is not fully supported by the HCD-MS/MS data, and in some cases, it does not agree with the predicted modification site (indicated by an asterisk).

[112-122]

EGF3: DVRACSSAPCANNGTCVSLDDGLYECSCAPGYSGKDCQ

**A** Reporter DLK1 expressed in wild-type HEK293 cells

DLK1 [112-122] Unglycosylated peptide ($m/z$ = 608.2424, $z$ = 2)

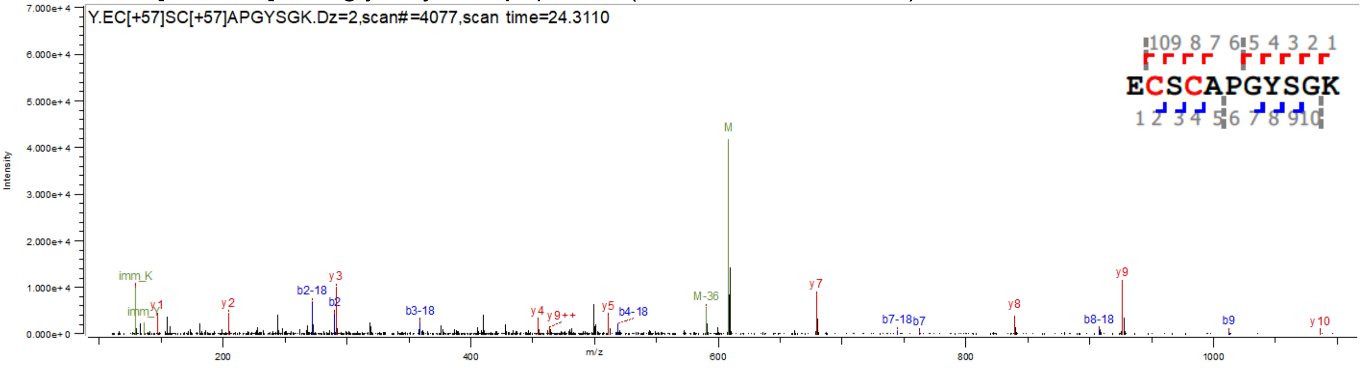


**B** Reporter DLK1 expressed in *POFUT1*-deficient HEK293 cells

DLK1 [112-122] Unglycosylated peptide ($m/z$ = 608.2425, $z$ = 2)

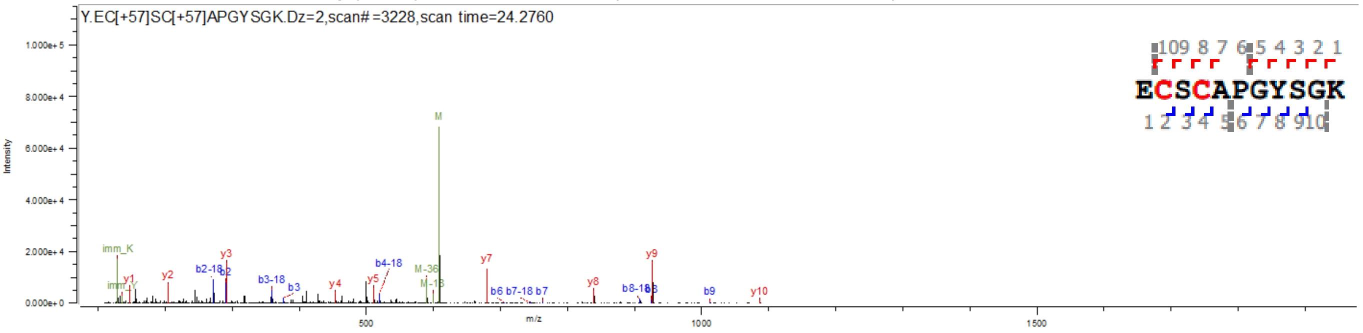


**C** Reporter DLK1 expressed in *POGLUT1*-deficient HEK293 cells

DLK1 [112-122] Unglycosylated peptide ($m/z$ = 608.2426, $z$ = 2)

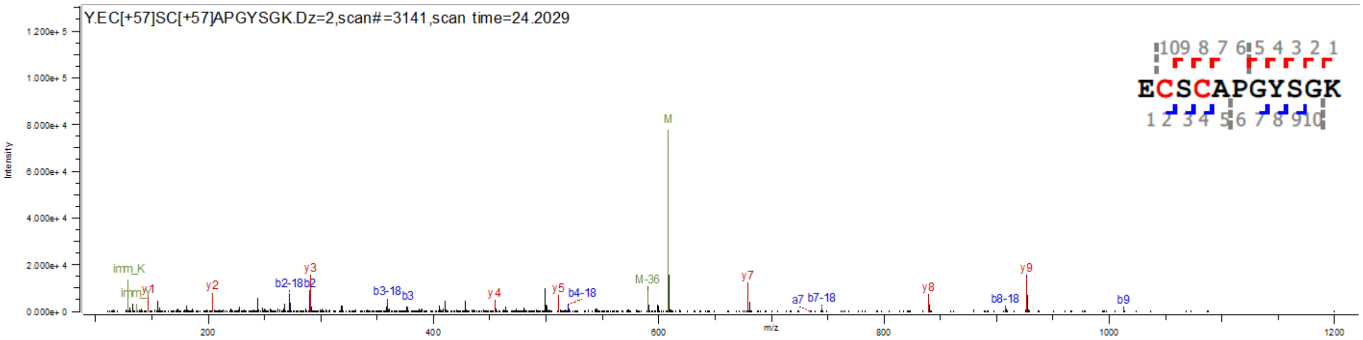


**Supporting Figure S14. MS/MS spectra of the DLK1[112-122] fragment of reporter DLK1.**
Byonic-assisted annotation of HCD-MS/MS spectra showing proteolytic fragments of DLK1[112-122]. Reporter DLK1 was prepared from wild-type HEK293 cells (A) or mutant cells lacking POGLUT1 (B) or POGLUT1 (C). After digestion with trypsin and chymotrypsin, the proteolytic fragments were treated with PNGase to remove N-glycans, and O-glycan modifications were analyzed by LC-MS/MS. The amino acid residue located at the putative O-GlcNAc modification site is indicated in green at the top of the figure.

Reporter DLK1 expressed in wild-type HEK293 cells

128-150

EGF4: KDGPCVI**N**GSPCQHGG**T**CVDDEGRASHASCLCPPGF**S**GNFCE

DLK1 [128-150]: Deamidation of Asn, dHex [1] (*m/z* = 878.6886, *z* = 3)

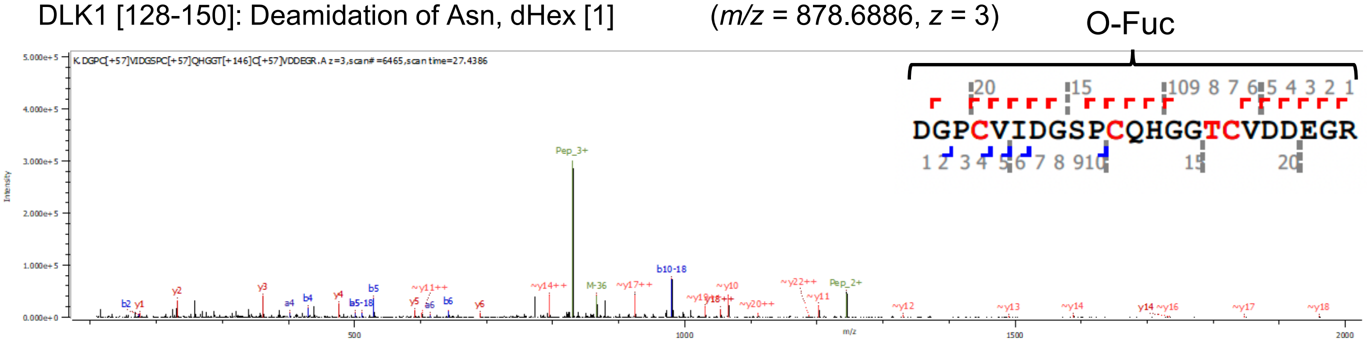


DLK1 [128-150]: dHex [1] (*m/z* = 878.3605, *z* = 3)

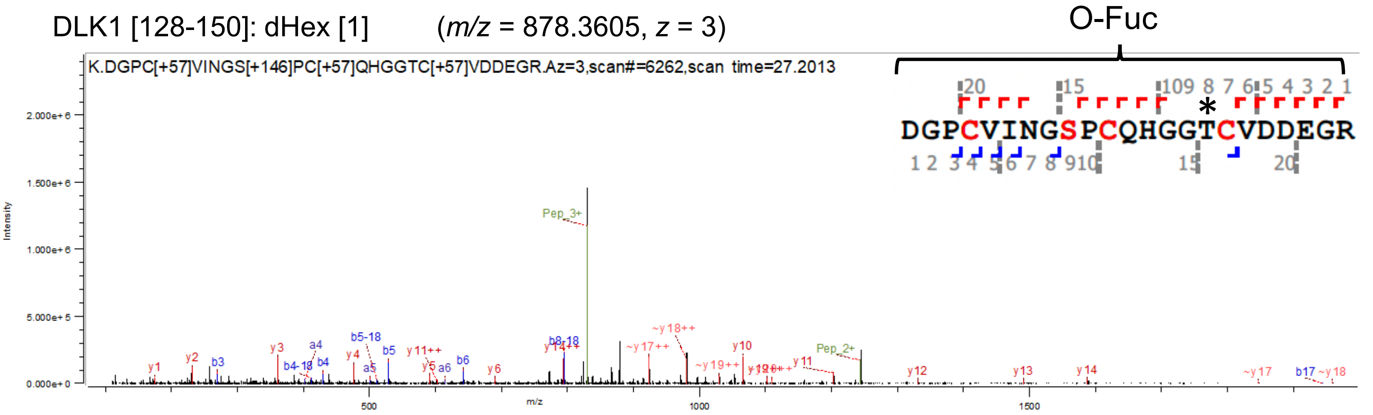


**Supporting Figure S15. MS/MS spectra of the DLK1[128-150] fragment of reporter DLK1 expressed in wild-type HEK293 cells.**

Byonic-assisted annotation of HCD-MS/MS spectra showing proteolytic fragments of DLK1[128-150] modified with N-glycans and/or the indicated O-glycans. Reporter DLK1 was prepared from transfected HEK293 cells and digested with trypsin and chymotrypsin. Proteolytic fragments were treated with PNGase to remove N-glycans. The resulting conversion of asparagine (N) to aspartic acid (D) (deamidation) and O-glycan modifications were analyzed using LC-MS/MS. Amino acid residues located at the putative modification sites for O-Fuc and N-glycan are indicated in red and brown, respectively, at the top of the figure. The modification site for the proposed O-glycan structure in the inset is not fully supported by the HCD-MS/MS data, and in some cases, it does not agree with the predicted modification site (indicated by an asterisk).

Reporter DLK1 expressed in *POFUT1*-deficient HEK293 cells

128-150

EGF4: KDGPCVINGSPCQHGGTCVDDEGRASHASCLCPPGFSGNFCE

DLK1 [128-150]: Deamidation of Asn (*m/z* = 830.0027, *z* = 3)

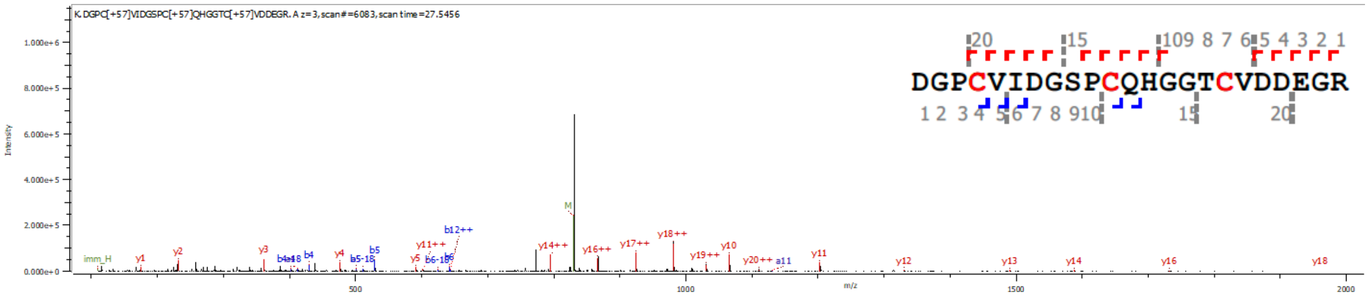


DLK1 [128-150]: Deamidation of Asn (*m/z* = 1244.0079, *z* = 2)

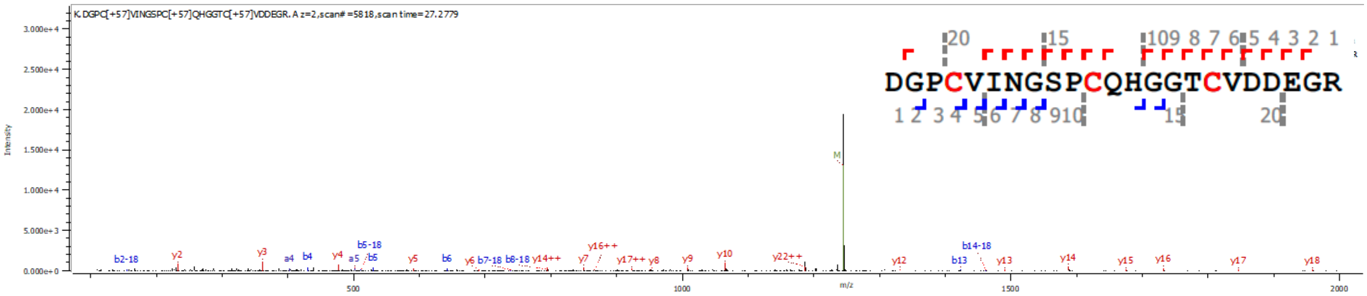


DLK1 [128-150]: Unglycosylated peptide (*m/z* = 829.6750, *z* = 3)

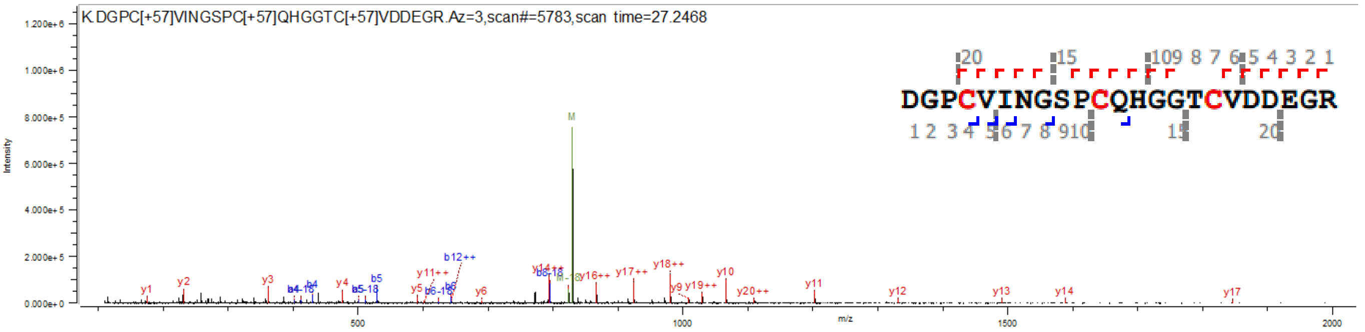


**Supporting Figure S16. MS/MS spectra of the DLK1[128-150] fragment of reporter DLK1 expressed in *POFUT1*-deficient HEK293 cells.**

Byonic-assisted annotation of HCD-MS/MS spectra showing proteolytic fragments of DLK1[128-150] modified with or without N-glycans. The reporter DLK1 was prepared from *POFUT1*-deficient HEK293 cells and digested with trypsin and chymotrypsin. Proteolytic fragments were treated with PNGase to remove N-glycans. The resulting conversion of asparagine (N) to aspartic acid (D) (deamidation) and O-glycan modifications were analyzed using LC-MS/MS. Amino acid residues located at the putative modification sites for O-Fuc and N-glycan are indicated in red and brown, respectively, at the top of the figure.

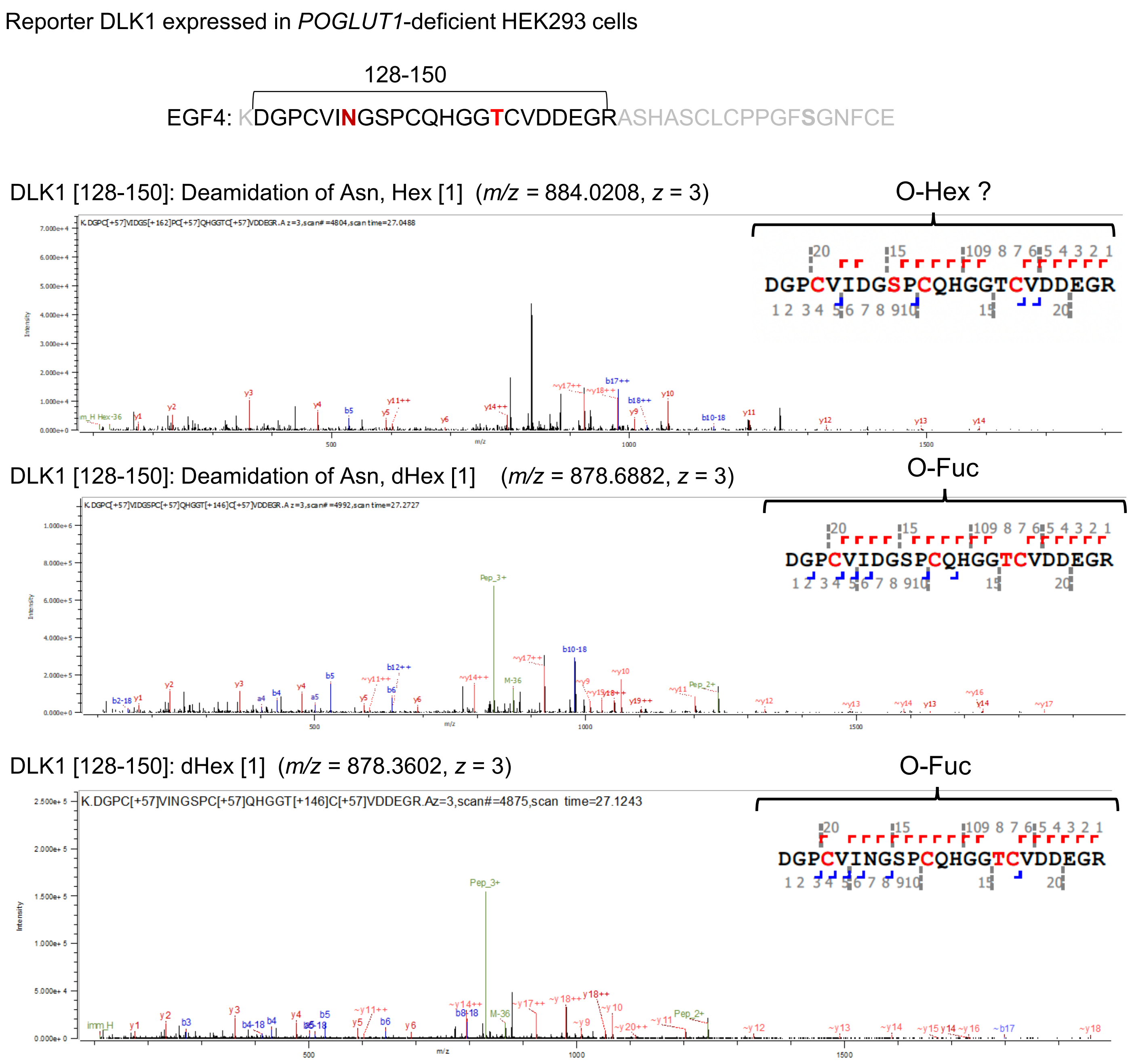


**Supporting Figure S17. MS/MS spectra of the DLK1[128-150] fragment of reporter DLK1 expressed in *POGLUT1*-deficient HEK293 cells.**

Byonic-assisted annotation of HCD-MS/MS spectra showing proteolytic fragments of DLK1[128-150] modified with N-glycans and/or indicated O-glycans. Reporter DLK1 was prepared from *POGLUT1*-deficient HEK293 cells and digested with trypsin and chymotrypsin. Proteolytic fragments were treated with PNGase to remove N-glycans. The resulting conversion of asparagine (N) to aspartic acid (D) (deamidation) and O-glycan modifications were analyzed using LC-MS/MS. The amino acid residues located at the putative modification sites for O-Fuc and N-glycan are indicated in red and brown, respectively, at the top of the figure. The modification site for the proposed O-glycan structure in the inset is not fully supported by the HCD-MS/MS data. The O-Hex modification site was not formally determined from the consensus sequence or MS/MS spectra.

Reporter DLK1 expressed in wild-type HEK293 cells

[151-166]

EGF4: KDGPCVINGSPCQHGGTCVDDEGRASHASCLCPPGFSGNFCE

DLK1 [151-166]: Unglycosylated peptide (*m/z* = 854.8644, *z* = 2)

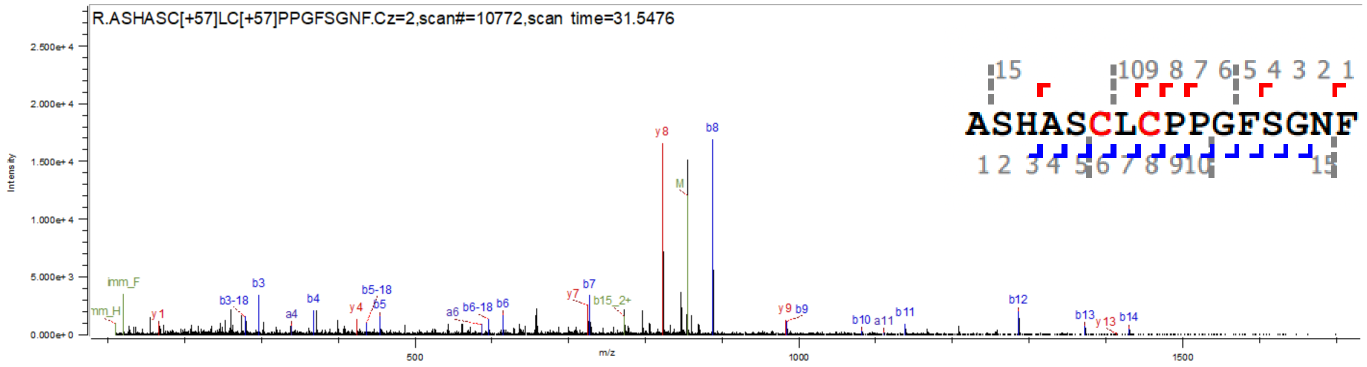


DLK1 [151-166]: Hex [1] (*m/z* = 935.8909, *z* = 2)

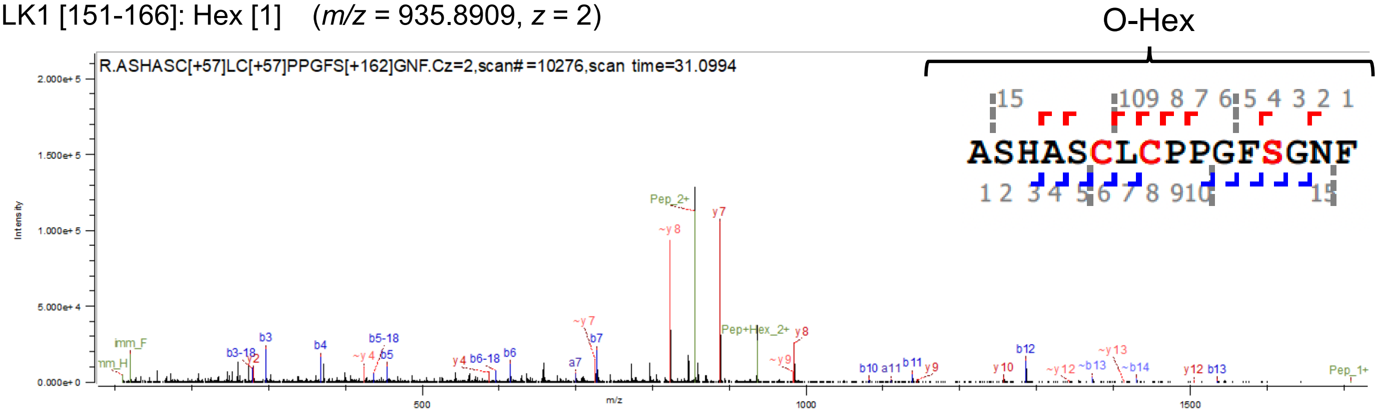


**Supporting Figure S18. MS/MS spectra of the DLK1[151-166] fragment of reporter DLK1 expressed in wild-type HEK293 cells.**
Byonic-assisted annotation of HCD-MS/MS spectra showing proteolytic fragments of DLK1[151-166] modified with or without O-Hex. Reporter DLK1 was prepared from transfected HEK293 cells and digested with trypsin and chymotrypsin. Proteolytic fragments were treated with PNGase to remove N-glycans, and O-glycan modifications were analyzed using LC-MS/MS. The amino acid residue located at the putative O-GlcNAc modification site is indicated in green at the top of the figure. The modification site for the proposed O-glycan structure in the inset is not fully supported by the HCD-MS/MS data.

Reporter DLK1 expressed in *POFUT1*-deficient HEK293 cells

[151-166]

EGF4: KDGPCVINGSPCQHGGTCVDDEGRASHASCLCPPGFSGNFCE

DLK1 [151-166]: Unglycosylated peptide (*m/z* = 854.8643, *z* = 2)

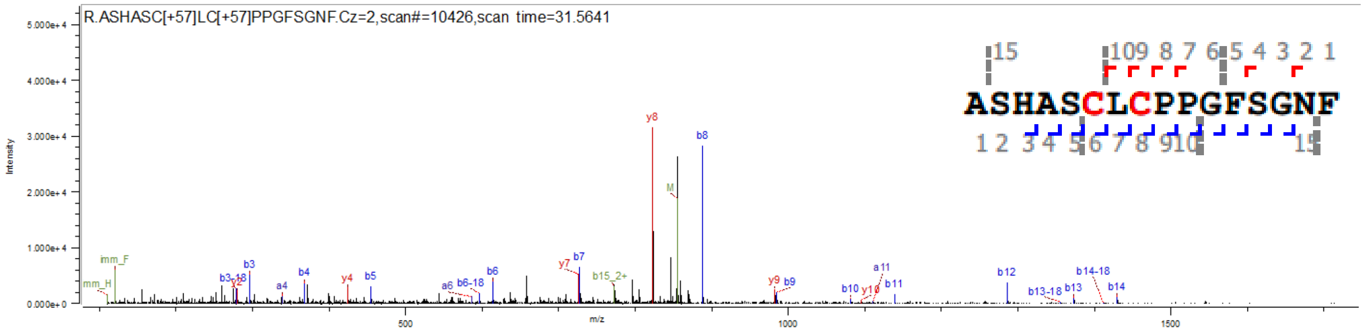


DLK1 [151-166]: Hex[1] (*m/z* = 935.8910, *z* = 2)

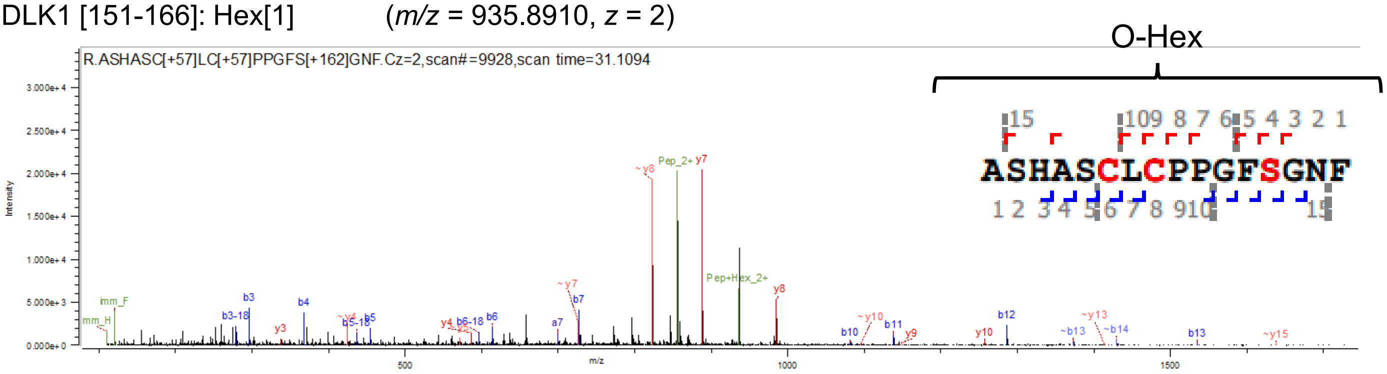


**Supporting Figure S19. MS/MS spectra of the DLK1[151-166] fragment of reporter DLK1 expressed in *POFUT1*-deficient HEK293 cells.**

Byonic-assisted annotation of HCD-MS/MS spectra showing proteolytic fragments of DLK1[151-166] modified with or without O-Hex. The reporter DLK1 was prepared from HEK293 cells and digested with trypsin and chymotrypsin. Proteolytic fragments were treated with PNGase to remove N-glycans, and O-glycan modifications were analyzed using LC-MS/MS. The amino acid residue located at the putative O-GlcNAc modification site is indicated in green at the top of the figure. The modification site for the proposed O-glycan structure in the inset is not fully supported by the HCD-MS/MS data.

Reporter DLK1 expressed in *POGLUT1*-deficient HEK293 cells

[151-166]

EGF4: KDGPCVINGSPCQHGGTCVDDEGRASHASCLCPPGFSGNFCE

DLK1 [151-166]: Unglycosylated peptide ($m/z$ = 854.8642, $z$ = 2)

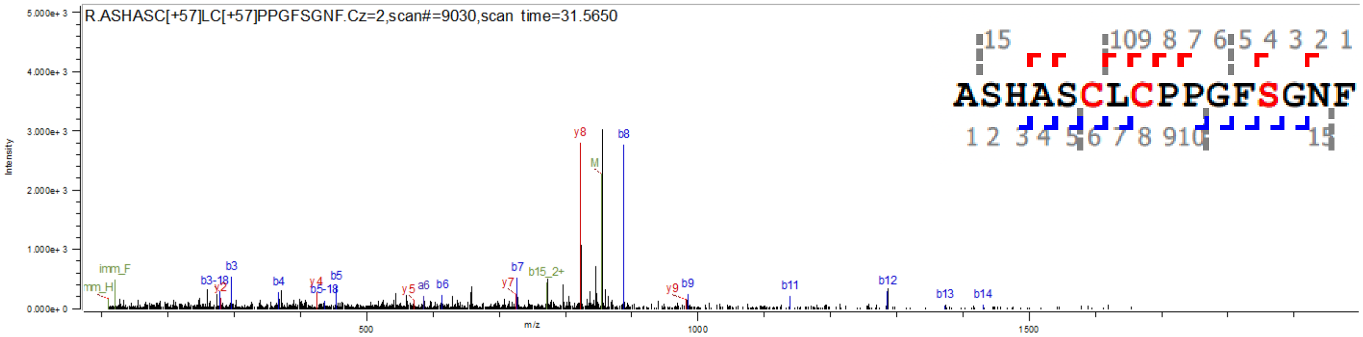


DLK1 [151-166]: Hex [1] ($m/z$ = 935.8909, $z$ = 2)

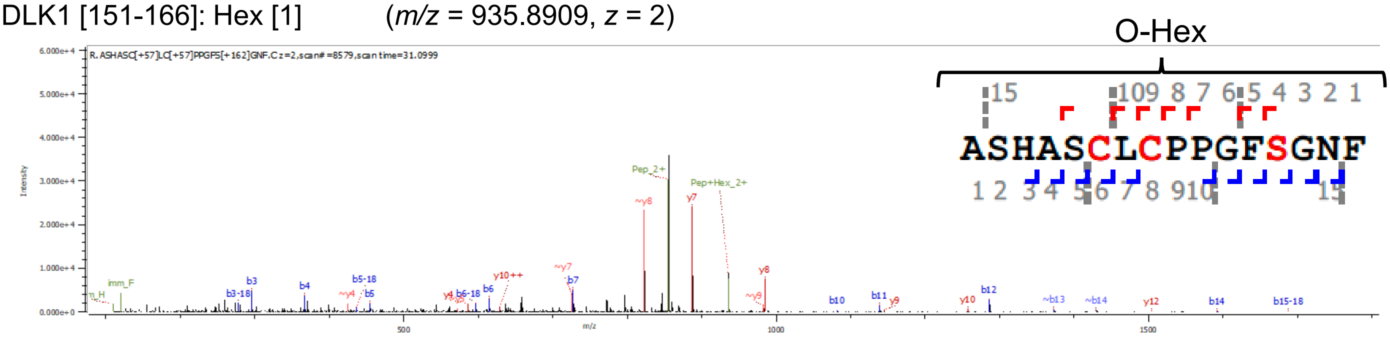


**Supporting Figure S20. MS/MS spectra of the DLK1[151-166] fragment of reporter DLK1 expressed in *POGLUT1*-deficient HEK293 cells.**

Byonic-assisted annotation of HCD-MS/MS spectra showing proteolytic fragments of DLK1[151-166] modified with or without O-Hex. The reporter DLK1 was prepared from HEK293 cells and digested with trypsin and chymotrypsin. Proteolytic fragments were treated with PNGase to remove N-glycans, and O-glycan modifications were analyzed using LC-MS/MS. The amino acid residue located at the putative O-GlcNAc modification site is indicated in green at the top of the figure. The modification site for the proposed O-glycan structure in the inset is not fully supported by the HCD-MS/MS data.

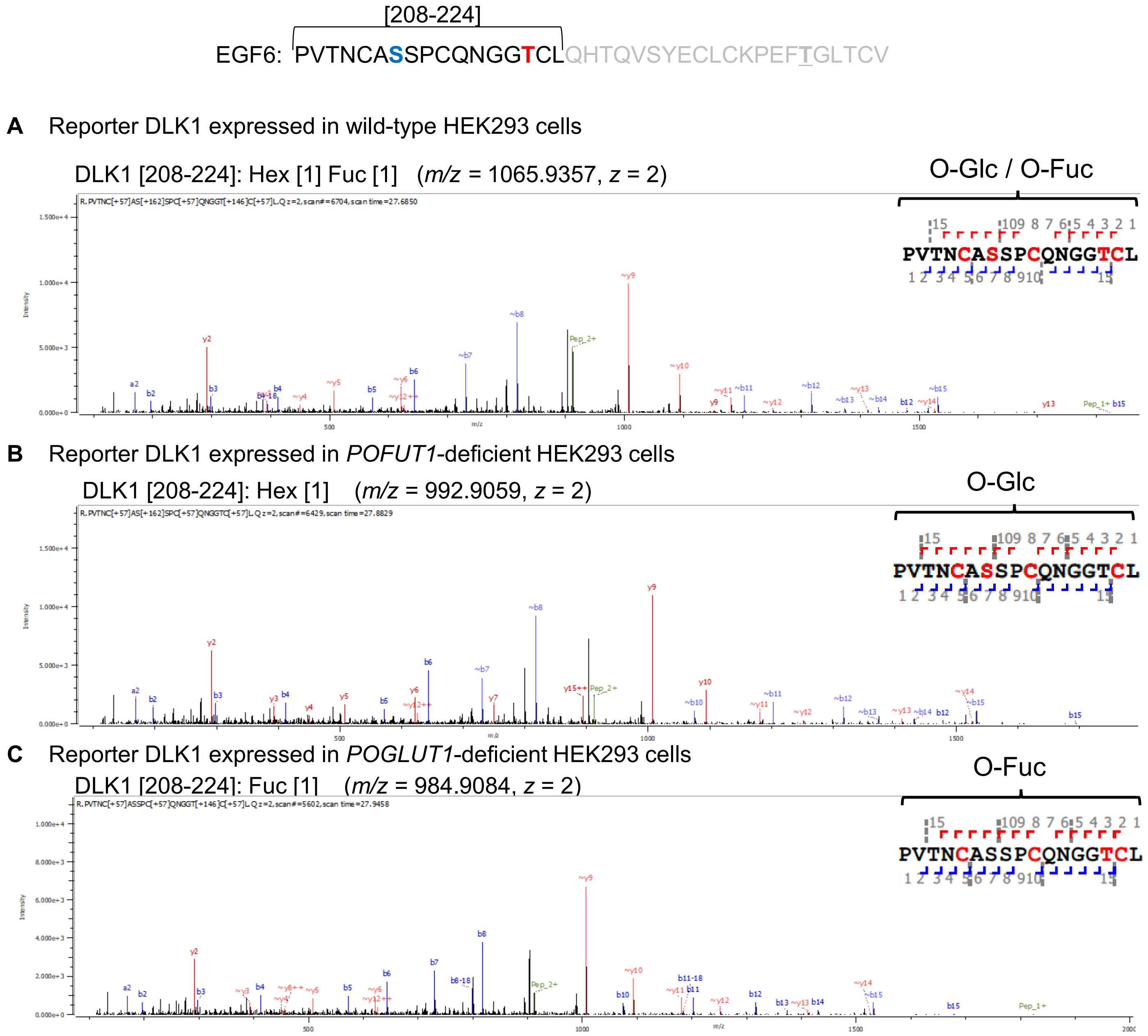


**Supporting Figure S21. MS/MS spectra of the DLK1[208-224] fragment of reporter DLK1.**
Byonic-assisted annotation of HCD-MS/MS spectra showing proteolytic fragments of DLK1[208-224]. The reporter DLK1 was prepared from wild-type HEK293 cells (A) or mutant cells lacking POGLUT1 (B) or POGLUT1 (C). After digestion with trypsin and chymotrypsin, the proteolytic fragments were treated with PNGase to remove N-glycans, and O-glycan modifications were analyzed using LC-MS/MS. The amino acid residues located at the putative modification sites for O-Glc and O-Fuc are indicated in green and red, respectively, at the top of the figure. The modification site for the proposed O-glycan structure in the inset is not fully supported by the HCD-MS/MS data.

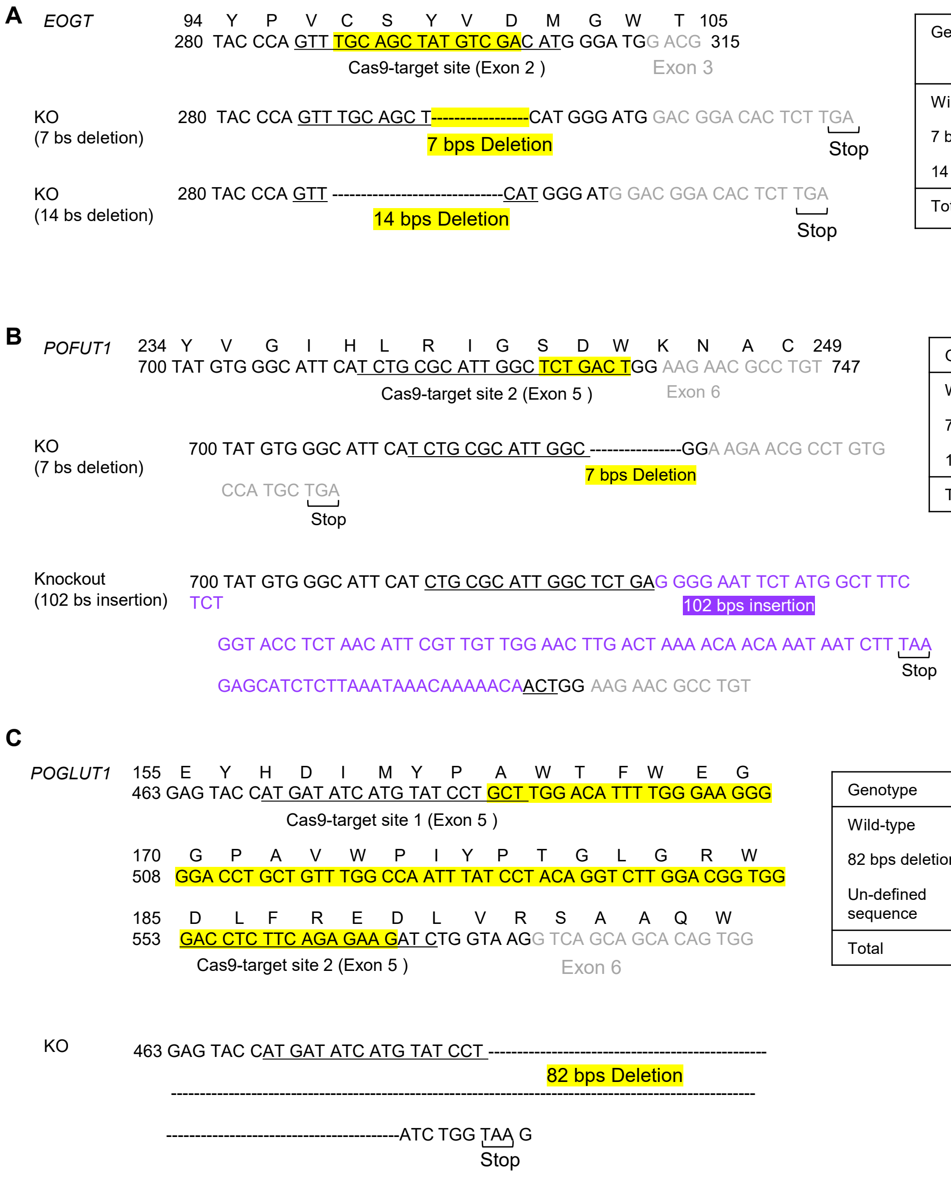


| Genotype | Number | | |
|---|---|---|---|
| | KO1 | KO2 | KO3 |
| Wild-type | 0 | 0 | 0 |
| 7 bps deletion | 6 | 7 | 6 |
| 14 bps deletion | 4 | 3 | 4 |
| Total | 10 | 10 | 10 |

| Genotype | Number |
|---|---|
| Wild-type | 0 |
| 7 bps deletion | 6 |
| 102 bps insertion | 4 |
| Total | 10 |

| Genotype | Number |
|---|---|
| Wild-type | 0 |
| 82 bps deletion | 7 |
| Un-defined sequence | 3 |
| Total | 10 |

**Supporting Figure 22. Deleted genomic regions in *EOGT, POFUT1, or POGLUT1* knockout cells derived from HEK293 cells.** Genomic regions deleted using the CRISPR/Cas 9 system are indicated in *EOGT*-deficient cells (A), *POFUT1*-deficient cells (B), and *POGLUT1*-deficient cells (C). The yellow maker indicates the deleted region. The inserted sequence is indicated in purple. Genome regions, including the Cas9-target site, were amplified using polymerase chain reaction-based TA cloning. The sequence of each clone was analyzed. The number of each genotype is shown in the table to the right.

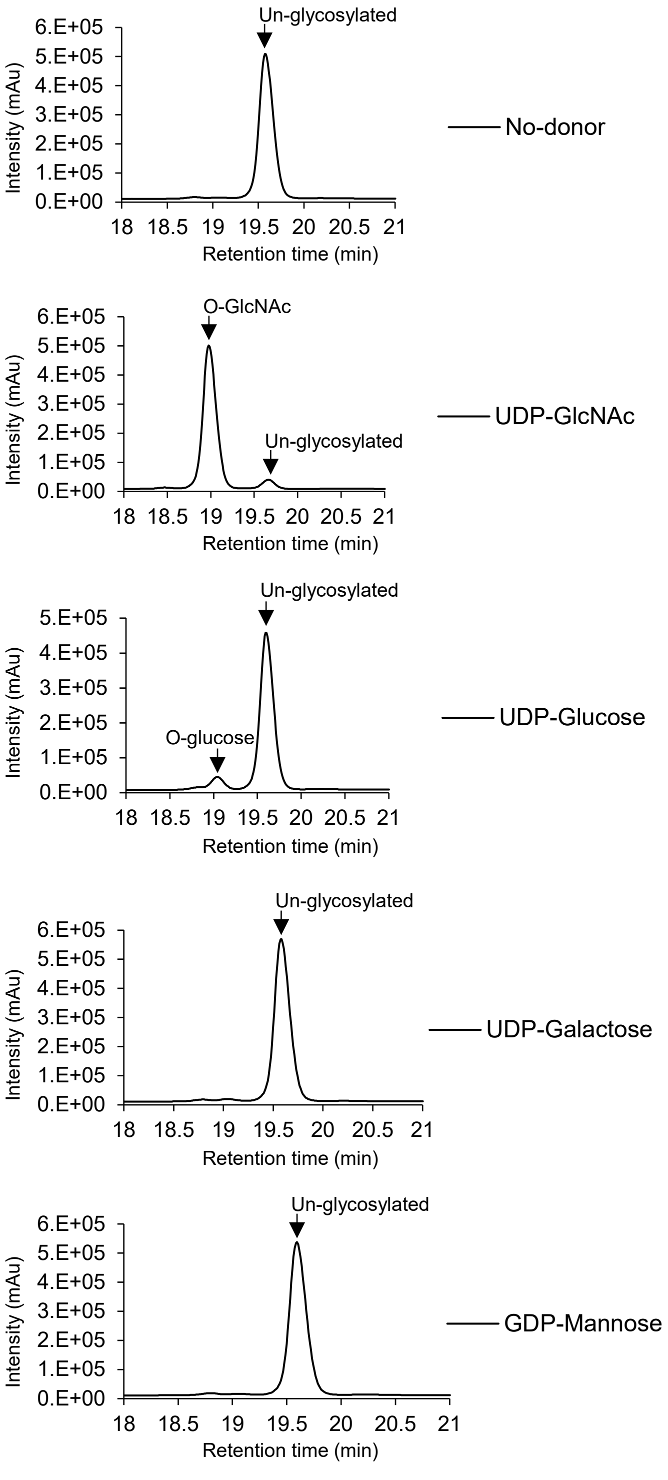


**Supporting Figure 23. *In vitro* EOGT enzyme assay.** EOGT expressed and purified from HEK293T cells was incubated with *Drosophila* NOTCH EGF20 (dEGF20) and the donor substrate as UDP-sugars indicated, at 37 ° C for 18 h. The reaction products were separated by reverse-phase high-performance liquid chromatography using a C18 column as previously described [1].

### Experimental Procedures for Supporting Figure 23.

FLAG-tagged EOGT was overexpressed in HEK293T cells and purified, as previously described (Ref. Tsukamoto et al, BBRC, 2024). Five-hundred nanograms of FLAG-tagged EOGT was incubated with 10 μg dEGF20 and 100 μM donor substrate in 125 mM HEPES-NaOH buffer (pH 7.0) with 10 mM $MgCl_2$ at 37 ° C for 18 h. The reaction volume was 50 μL. The reaction products were diluted to 1 mL and separated by reverse-phase high-performance liquid chromatography using a C18 column as previously described [1].
